# Greener, Wilder, Better: Valuing Complexity and Diversity in Urban Riparian Areas

**DOI:** 10.1101/2025.10.24.684068

**Authors:** Arpad Erik Thoma, Henrique M. Pereira, Marek Giergiczny

## Abstract

(1) Riparian areas are among the most valuable ecosystems on earth due to their high biodiversity and contributions to human well-being, yet urban riparian areas face significant threats from urbanization and associated expansion. Current management strategies do not consider all the contributions of urban green spaces to human well-being. Over the last decade stated preference studies have been widely used to gain insights into public attitudes towards urban green spaces. However, whether and to what extent the public values urban biodiversity in urban riparian areas remains unclear.
(2) We used a discrete choice experiment approach to determine public preference for urban biodiversity and management interventions in urban riparian areas through a discrete choice experiment conducted in Poland.
(3) We found strong positive preferences for highly complex and dense vegetation along urban riparian areas, indicating a public inclination towards enhancing natural biodiversity in public spaces. Additionally, respondents expressed a preference for reducing impermeable surfaces, such as concrete pathways, highlighting the value placed on minimizing urban encroachment.
(4) Willingness-to-pay (WTP) results suggested that the public in Poland is prepared to financially support certain biodiversity conservation measures and land-conversion interventions in urban green spaces.
(5) These results underscore the importance of integrating ecological research and public preferences into urban riparian management strategies, emphasizing the need for policy approaches that prioritize biodiversity conservation while balancing urban development.

## Introduction

Sustainable planning and management of urban futures face immense challenges to ensure progress in social and economic development as well as the protection and preservation of the environment (United Nations, 2019). Today, more than half of the world’s population lives in urban areas, and this share is projected to rise to nearly 70% by 2050. This rapid urbanization comes at a significant ecological cost - natural habitats are being fragmented, ecosystem functions disrupted, and biodiversity increasingly threatened (IPBES, 2019; McDonald et al., 2020; Simkin et al., 2022). In addition to the immediate consequences of urban expansion, its indirect effects are escalating. While consuming most of the world’s resources, cities contribute over 70% of the worldwide CO2 emissions and produce half of the world’s waste (Lucertini & Musco, 2020). The resulting environmental pressure affects natural areas and biodiversity even far from cities (Mansur et al., 2022).

Preserving urban biodiversity is therefore a critical component of sustainable cities. It enhances ecosystem resilience to climate change (Ratzke, 2023) and supports human-nature connections essential for fostering environmental stewardship (Miller, 2005). Yet, effective conservation within urban landscapes requires a better understanding of how urban ecosystems are valued by the public and how management interventions are perceived.

Urban green spaces (UGS) - including parks, forests, rivers, and wetlands - are increasingly recognized for their multifunctional role in delivering both ecological and social benefits (Carrus et al., 2015; Zhang et al., 2020). Among these, urban riparian areas occupy a unique position. Situated at the junction between land and water, they provide important habitat corridors, support high levels of biodiversity, and offer valuable recreational and cultural services (Groffman et al., 2003). However, riparian ecosystems in cities are often heavily altered – rivers have been channelized, paved over, or simplified and adjacent areas have been transformed into impermeable surfaces - leading to a loss of ecological function including water filtration, habitat provision, and thermoregulation of urban microclimates (Peroni et al., 2023).

Despite their value, management strategies of urban green and blue spaces remain grounded in standardized criteria that overlook the full range of their contributions to human well-being (Reyes-Riveros et al., 2021) and often fail to incorporate ecological evidence (Lepczyk et al., 2017). Moreover, planning decisions rarely reflect the diversity of public preferences, limiting both the social relevance and ecological effectiveness of management interventions.

Valuation studies can help bridge this gap. Stated preference methods, particularly discrete choice experiments (DCEs), have proven useful in capturing how individuals value non-market benefits of urban biodiversity and are willing to pay (WTP) for specific improvements (Champ et al., 2017; Hanley et al., 1998). DCEs allow researchers to disentangle preferences for multiple attributes of ecosystems, offering insights that can guide targeted and publicly supported management strategies. However, in its complexity as an ecological concept, biodiversity cannot be captured directly for valuation purposes, leading to the use of bridging concepts or proxys like naturalness or even nature. Findings from such studies are often contradictory. For instance, a study by Stessens et al. (2020) found that naturalness was perceived as less important compared to other qualities of urban green spaces, while a study by Bronnmann et al. (2023) found a positive mean WTP for an increase of naturalness of UGS. To what extent the public values urban biodiversity and are willing to pay for its preservation remains unclear (Ratzke, 2023).

Urban blue infrastructure, including rivers, canals, and adjacent green corridors, has also received less valuation-attention than other UGS types (Bockarjova et al., 2020) - especially in Central and Eastern Europe, where rapid urban transformation coincides with high biodiversity potential and distinctive socio-cultural relationships with nature (Noszczyk et al., 2023).

This paper addresses these gaps by focusing on urban riparian areas as multifunctional spaces where ecological processes and human use intersect, yet where management rarely accounts for the public’s specific preferences for ecological features. We advance existing valuation research by systematically decomposing biodiversity into ecologically grounded, directly observable attributes - including vegetation characteristics, landscape complexity, vegetation cover, species richness, presence of dead wood, management intensity, and the extent of ground sealing. Using a discrete choice experiment in Polish cities, we quantify the relative importance of these attributes and translate preferences into WTP estimates. This multidimensional, attribute-based approach allows us to disentangle how the public values distinct components of riparian biodiversity and to provide actionable, monetized evidence that can guide urban river management strategies which are ecologically meaningful, socially supported, and financially feasible.

## 2. Methods

### 2.1 Study area

Our study focuses on the valuation of urban riparian areas in Poland, with particular emphasis on how residents of cities with prominent blue infrastructure perceive and value different biodiversity and management attributes of these spaces. Poland has been undergoing a steady urban transformation, rising from around 55% of its total population in 1975 (34 million residents) to nearly 60% of its 38 million residents living in urban areas today. (Local Data Bank, 2022). Major urban centers - such as Warsaw, Kraków, Łódź, Poznań, and Wrocław - exemplify rapidly developing cities where pressures on land use, biodiversity, and public space are intensifying (Noszczyk et al., 2023). We targeted adult residents living in cities with more than 20,000 inhabitants and significant river infrastructure, defined by the presence of major rivers such as the Wisła (Vistula), Odra (Oder), and Warta (Figure 1). These rivers traverse dense urban fabrics and are often heavily engineered - shaped by historical modifications, flood control measures, and ongoing urban redevelopment. Nevertheless, they retain pockets of ecological potential and serve as vital spatial anchors for urban blue-green infrastructure. The sample of our study included residents from cities of varying sizes, geographic locations, and riparian landscapes, ensuring broad representation across regional and socio-demographic context.

**Figure 1:**
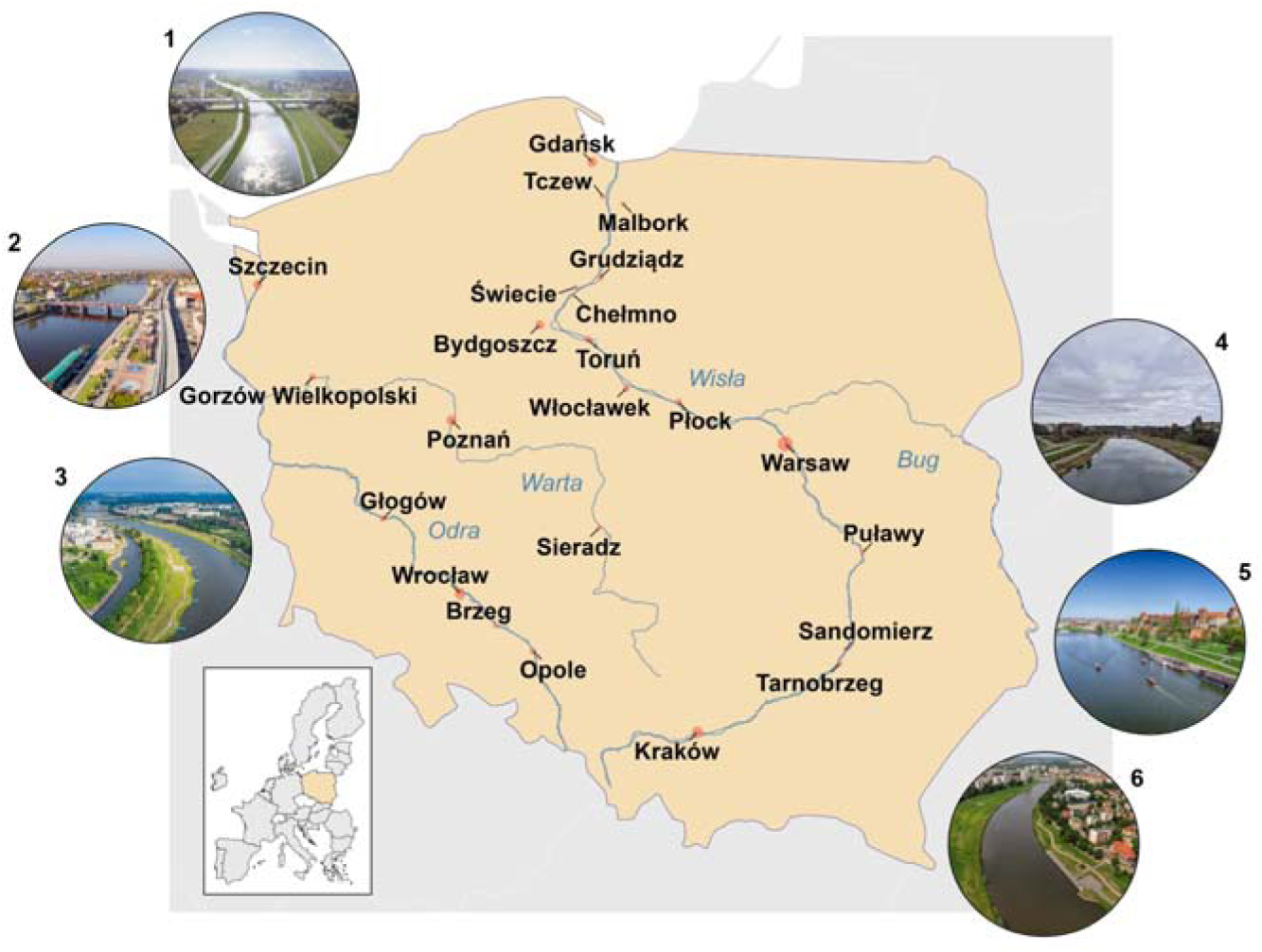
Map of Poland indicating the study area. Displayed are cities with > 20,000 inhabitants along significant river infrastructure. Red circle sizes at city locations indicate the different population sizes. The four main rivers of Poland are displayed in blue and labeled accordingly. Images 1-6 represent selected photographs of urban riparian areas along the Warta river in (1) Poznań, along the Odra river in (2) Gorzów Wielkopolski and (3) Wrocław and along the Wisła river in (4) Sieradz, (5) Kraków and (6) Opole.

### 2.2 Identification of urban riparian attributes and levels

To explore public preference for urban riparian areas, we designed a discrete choice experiment (DCE) grounded in attributes that reflect meaningful and measurable outcomes of urban riparian management. Building on ecological literature (Groffman et al., 2003; Naiman et al., 1993; Naiman et al., 2005; Peroni et al., 2023), expert input, and adaptation of the QBR (Qualitat del Bosc de Ribera) index (Munné et al., 2003), we focused on features that could plausibly result from nature-based restoration or shifts in management practices. To incorporate the complexity of the concept biodiversity in our study we followed the suggestions of Bartkowski et al. (2015) and selected a variety of attributes describing different aspects of biodiversity, rather than relying on bridging concepts. We put considerable effort into the selection of attributes, to make sure the attributes could be observed and understood by laypeople. The design aimed to balance ecological relevance with public salience, enabling respondents to weigh trade-offs between biodiversity enhancement, urban development, and management intensity.

After considering an initial list of ten attributes, we retained eight that captured key dimensions of urban riparian areas (Table 1). Together, these attributes provide a multidimensional framework for evaluating public support for urban riparian interventions. These were grouped into three categories: (1) Vegetation characteristics, (2) Species diversity and microhabitat provision and (3) Human intervention and land conversion.

**Table 1:**
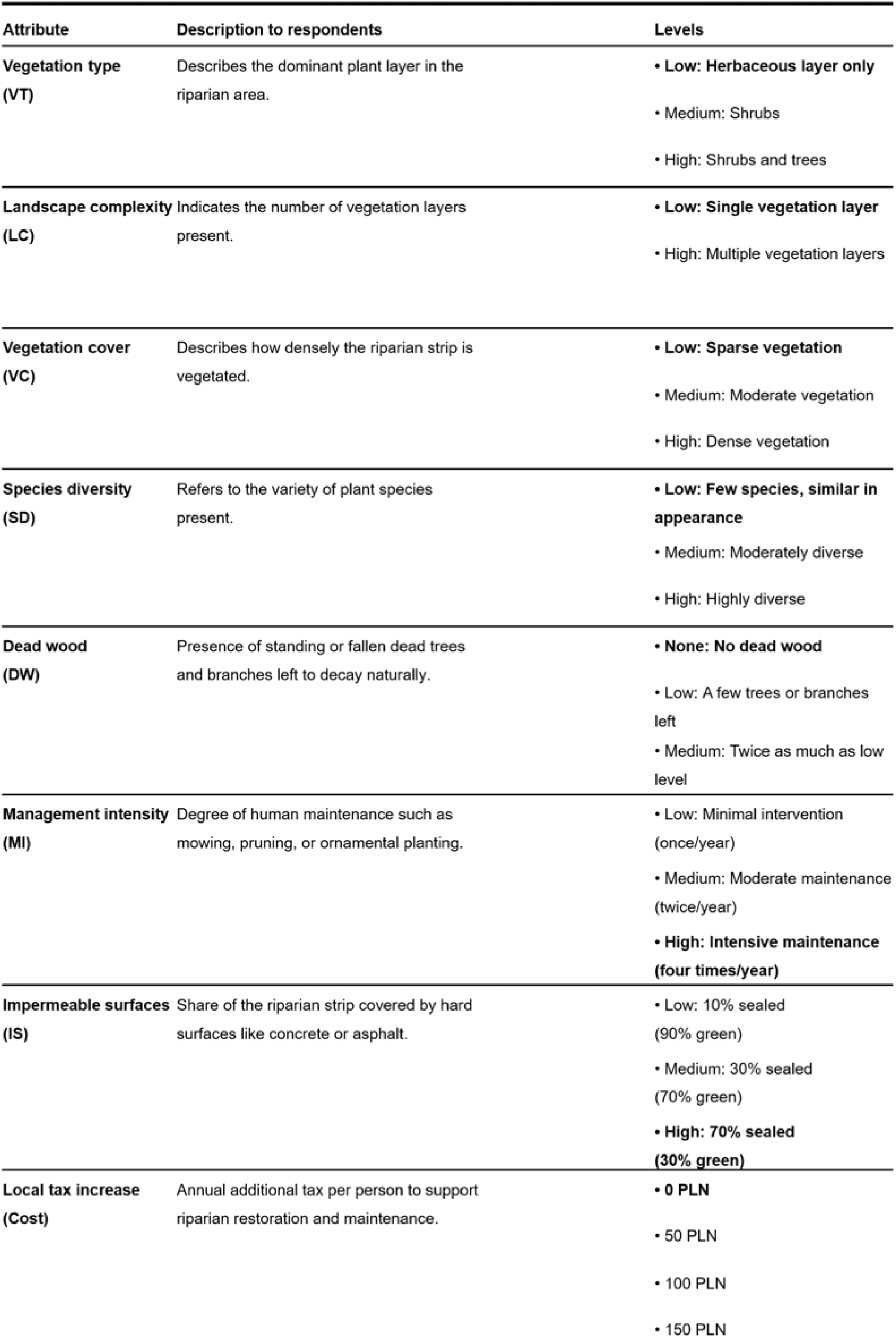
Overview of the attributes and their levels used in the DCE. Status quo levels are shown in bold.

**Vegetation characteristics**, the first set of attributes within our framework, addressed the structural complexity of riparian ecosystems (Table 1). The attribute ‘vegetation type’ referred to the main vegetation layer present, varying between herbaceous layer, shrubs, and trees, while ‘landscape complexity’ indicated the presence of either a single vegetation type or a mix of different vegetation types. The ‘vegetation cover’ attribute assessed the total amount of plant cover within the riparian area and serves as an indicator for connectivity, addressing habitat fragmentation that has led to significant obstacles to ecosystem functioning within urban riparian areas.

**Species diversity and microhabitat provision**, the second element of our attribute framework, focused on the biological richness and ecological complexity within riparian systems (Table 1). ‘Species diversity’ referred specifically to the different plant species within the various vegetation types, while ‘dead wood’ addressed the presence or absence of standing dying trees or laying dead wood along the riparian area. This attribute presented a crucial biodiversity indicator, referring to microhabitats for insects and microbes, as well as nutrient provision, allowing natural processes such as decomposition to occur freely within the ecosystem.

**Human intervention and land conversion**, the third element, concerned the degree of human intervention and urban development pressure (Table 1). The attribute ‘management intensity’ refers to the amount of human-induced changes due to activities such as mowing, pruning and planting of ornamental plants, with levels varying on a scale from intensive management to completely hands-off management, although this approach may also entail negative perceived consequences for people’s interests regarding maintenance and aesthetics. The attribute ‘impermeable surfaces’ referred to the share of concrete sealed areas along the urban riparian area, addressing urban expansion and subsequent habitat degradation. The levels for this attribute ranged from increased intensification of sealed surfaces to reduced impermeability.

To ensure accessibility, we employed plain language and avoided technical jargon. Each attribute was visualized using Twinmotion version 2023 (Twinmotion, 2023), which enabled the creation of realistic depictions of vegetation types, habitat features, and land use patterns familiar within a Central European context (Figure 2). Such visualizations have been shown to improve comprehension and engagement in discrete choice experiments (Eberhard, 2023; Shr et al., 2019), supporting their use in our approach. The use of the software allowed us to keep the attributes levels constant or to change them, while always using the same background and light settings. We produced a total of 119 visualizations for this study, each being unique in the combination of attributes and their levels.

**Figure 2:**
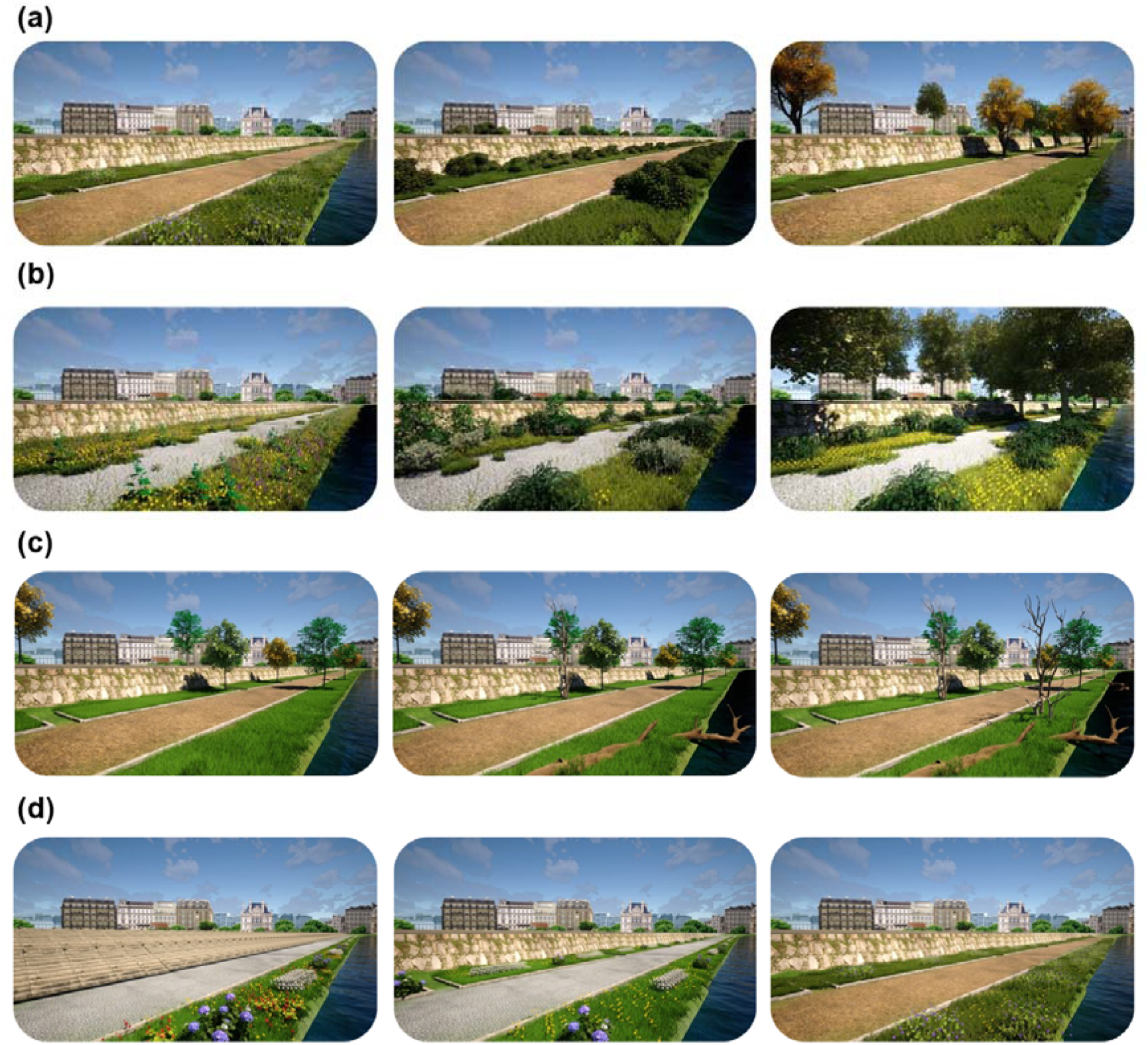
Example visualizations of urban riparian areas using the Twinmotion software. Row (a) shows urban riparian areas with changing vegetation type and a constant level of management intensity, starting with a low level (herbaceous layer only) on the left, a medium level (shrub layer) of vegetation type in the middle and a high level (trees) on the right. Row (b) shows landscapes with changing landscape complexity and a constant level of management intensity, starting with a low level of landscape complexity on the left and in the middle and a high level of landscape complexity on the right (presence of shrubs and trees simultaneously). Row (c) shows urban riparian areas with changing amount of standing or laying deadwood and constant levels of species diversity and management intensity, starting with no dead wood present on the left, low amount of deadwood in the middle and medium amount of dead wood on the right. Row (d) shows urban riparian areas with changing share of impermeable surfaces compared to vegetated area, starting with a share of 70% impermeable surface on the left, 30% impermeable surface in the middle and 10% impermeable surface on the right.

### 2.3 Choice experiment set-up

We used a discrete choice experiment (DCE) to assess public preferences and willingness to pay (WTP) for biodiversity-enhancing interventions in urban riparian areas in Poland. The survey was conducted online (CAWI mode) by a professional polling company, using stratified sampling to ensure representativeness across age, gender, region, and urban size. Only respondents living in cities with more than 20,000 inhabitants that have adjacent riparian areas were included.

The questionnaire comprised five sections (Supporting Information, S1). After collecting socio-demographic data, respondents were asked about their engagement with riparian areas, including frequency of visits and general environmental attitudes. The following section introduced the DCE attributes. These were explained in accessible language, accompanied by visual representations of each attribute level to enhance understanding. To promote reflection, respondents were also asked to rank attribute levels prior to completing the choice tasks.

The fourth section presented the hypothetical management program. Respondents were informed that changes to urban riparian areas would be funded through an annual increase in local taxes, which served as the payment vehicle in the DCE. To enhance incentive compatibility (Carson & Groves, 2007; Vossler & Watson, 2013) and reduce the risk of hypothetical bias (Champ et al., 2017; Kling et al., 2012), the scenario specified that, if the proposed program were implemented, the tax increase would be mandatory for all local residents, with no possibility to opt out. This follows best practice in stated preference valuation, where binding and unavoidable payment vehicles are recommended to mimic the conditions of a real economic decision and discourage free-riding (Carson & Groves, 2007; Champ et al., 2017; Johnston et al., 2017). By clearly stating the compulsory nature of the payment, respondents are more likely to treat the choice task as consequential, aligning their stated preferences with the true trade-offs they would face in reality.

Each respondent was presented with 16 choice cards (Figure 3), each displaying three alternatives: two hypothetical riparian management scenarios and one status quo option. Including a status quo alternative mirrors real-world decision-making contexts and provides a welfare-theoretic baseline necessary for consistent estimation of welfare measures (Louviere et al., 2000). The design was generated using a Bayesian efficient approach in Ngene 1.4 (ChoiceMetrics, 2024). It comprised 48 unique choice scenarios, divided into three blocks of 16 cards. A pilot survey (n = 100) informed the prior parameter values. To reduce ordering bias, the sequence of choice cards was randomized across respondents. Each choice set included five non-monetary attributes: vegetation type, landscape complexity, vegetation cover, species diversity, and management intensity. To reduce the complexity of the choice experiment—and thereby the cognitive burden from too many attributes—we presented the presence of dead wood and the proportion of impermeable surfaces selectively, on eight of the 16 choice cards; all other attributes appeared on all 16 cards.

**Figure 3:**
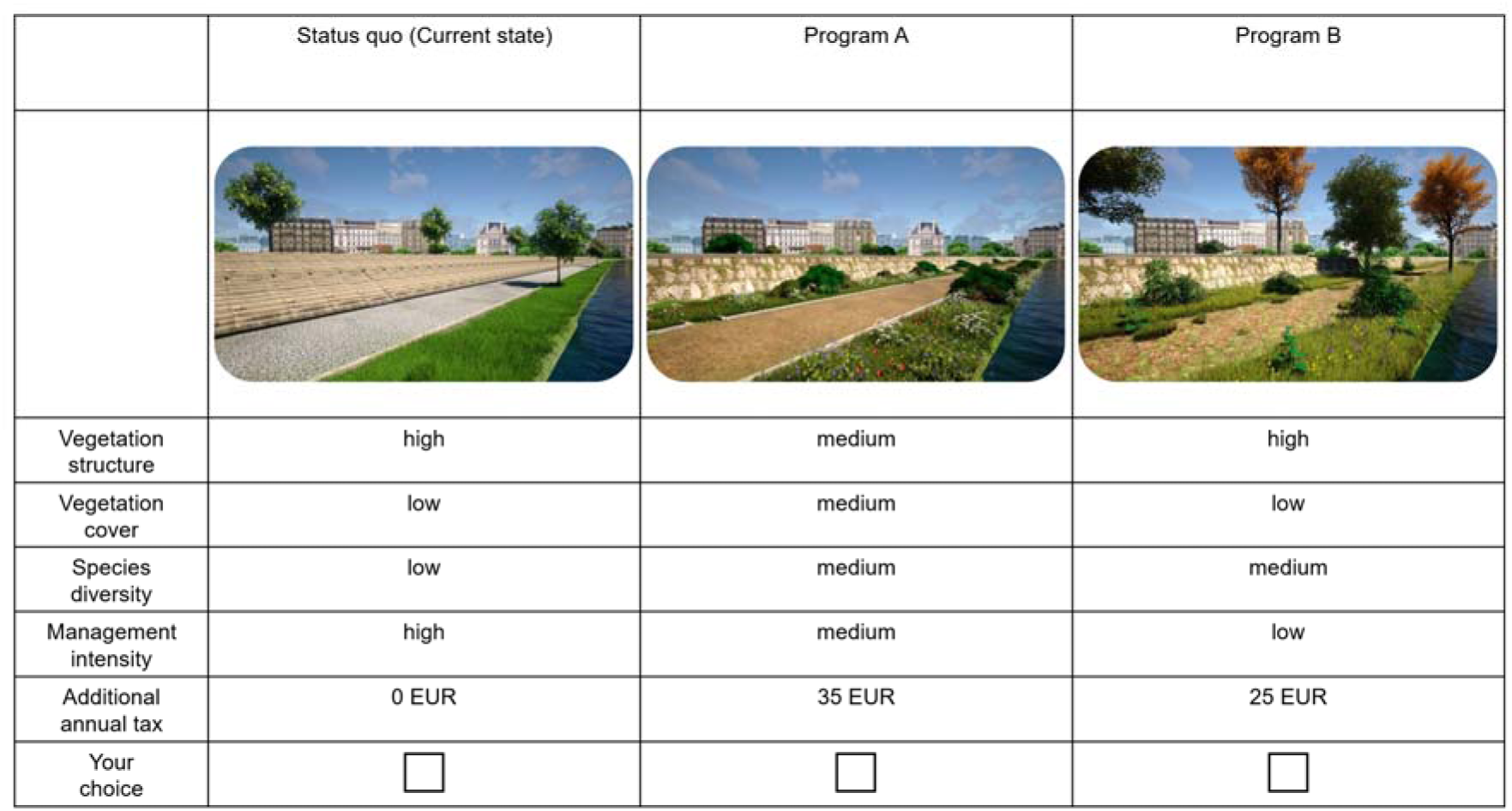
Example choice card.

All materials - including attribute texts and visualizations - were developed iteratively through focus group discussions and expert reviews. The content was designed to reflect ecologically meaningful changes while remaining comprehensible to the general public. The questionnaire was developed in English and translated into Polish by native speakers, with particular attention to linguistic clarity and accessibility. Ethical approval for this study was obtained from the Ethics Committee of the Faculty of Economic Sciences at the University of Warsaw. All collected data were anonymized and ensured that respondents were over the age of 18. Even though each respondent was only identifiable via a unique code (letters and numbers), this information was not used for this study. All respondents took part in this study voluntarily, and by submitting their answers, gave consent to use the anonymized data.

### 2.4 Choice experiment analysis

We applied a Mixed logit model (MXL) to analyze the data from the DCE. The model is rooted in random utility theory (McFadden, 1974), which assumes that an individual’s (*n*) preference can be decomposed into a deterministic component (*V_itn_*) and an unobservable, stochastic component (*ε_itn_*). This leads to the usual formulation of the utility that an individual derives from choosing alternative *i* at the choice occasion *t*,

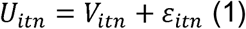

In this study, we specify the deterministic component of the utility as

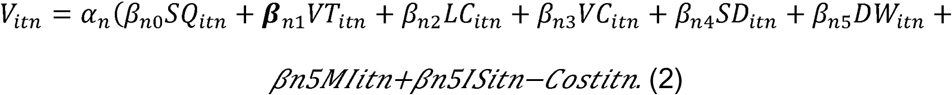

Here, the model in (2) is specified in the WTP-space (Train & Weeks, 2005), so that *α_i_* coefficient represents the marginal utility of money (confounded with the scale), whereas the *β* coefficients can be interpreted directly in monetary terms as a willingness to pay (WTP).

*SQ_itn_* is a dummy variable equal to 1 for the status quo (SQ) alternative, making *β_i_*_0_ the status quo alternative-specific constant (ASC), *VT_itn_* vegetation type, *LC_itn_* landscape complexity, *VC_itn_* vegetation cover, *SD_itn_* species diversity, *DW_itn_* the presence of dead wood, *IS_itn_* the proportion of impermeable surfaces. All non-monetary attributes were specified as categorical variables. Finnaly, *Cost_itn_* denotes the monetary attribute, expressed as the annual increase in local taxes and was coded as continuous variable.

We assume that coefficients that can be interpreted as WTP (*β_nj_*), follow a normal distribution. The marginal utility of money (*α_i_*) is assumed to follow a log-normal distribution.

Due to the unobserved stochastic terms, the likelihood function does not have an analytical form but instead is a multidimensional integral (3).

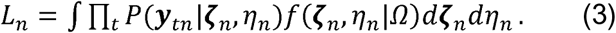

In (3), *f*(*ζ_n_*, *η_n_*|*Ω*) is a density function of the multivariate normal distribution, with the mean zero, and covariance matrix, *Ω*, being fully estimated. *P*(***y****_tn_*|*ζ_n_*, *η_n_*) denotes a choice probability with ***y****_tn_* being a vector of zeros and ones, with one indicating a chosen alternative. Assuming Gumbel-distributed error terms, *ε_itn_*, in (1), leads to a well-known multinomial logit choice probability,

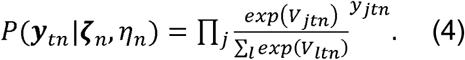

We estimated the model using the Maximum Simulated Likelihood approach with 1,000 scrambled Sobol draws to approximate the integral in (4).

In addition to the discrete choice experiment, we conducted a direct preference assessment to validate and contextualize the DCE findings. In this supplementary exercise, respondents were asked to identify their most and least preferred levels for each attribute independently. This approach provides a transparent and cognitively simpler assessment of preferences, helping to verify whether the preference patterns observed in the DCE align with respondents’ directly stated preferences.

## 3. Results

### 3.1 Respondent profile and recreational use of urban rivers

The final sample included 796 respondents living in towns and cities across Poland with populations above 20,000. The sample was structured to be broadly representative of this urban population in terms of age, gender, municipality size, and region. The main socio-demographic characteristics are summarized in Figure 4a. Urban rivers were used for a variety of activities (Figure 4b). The most common were leisure walking (91%) and landscape observation (46%), followed by cycling (40%) and nature observations (24%). Less frequent but notable activities included picnicking (22%), jogging (15%), and sunbathing (14%). Water-based activities such as swimming (5%), kayaking (3%), and boating (5%) were relatively uncommon. Respondents highlighted a mix of ecological and recreational factors as important to their well-being during river visits (Figure 4c). The most frequently mentioned were attractive landscapes (65%), natural vegetation (48%), walking paths (46%), and peace and quiet (40%). Water and air quality were also valued by a third of respondents. In contrast, infrastructure like restaurants/bars and cafés (7%) or recreational facilities (17%) were less frequently reported, indicating a general preference for nature-based over built features.

**Figure 4:**
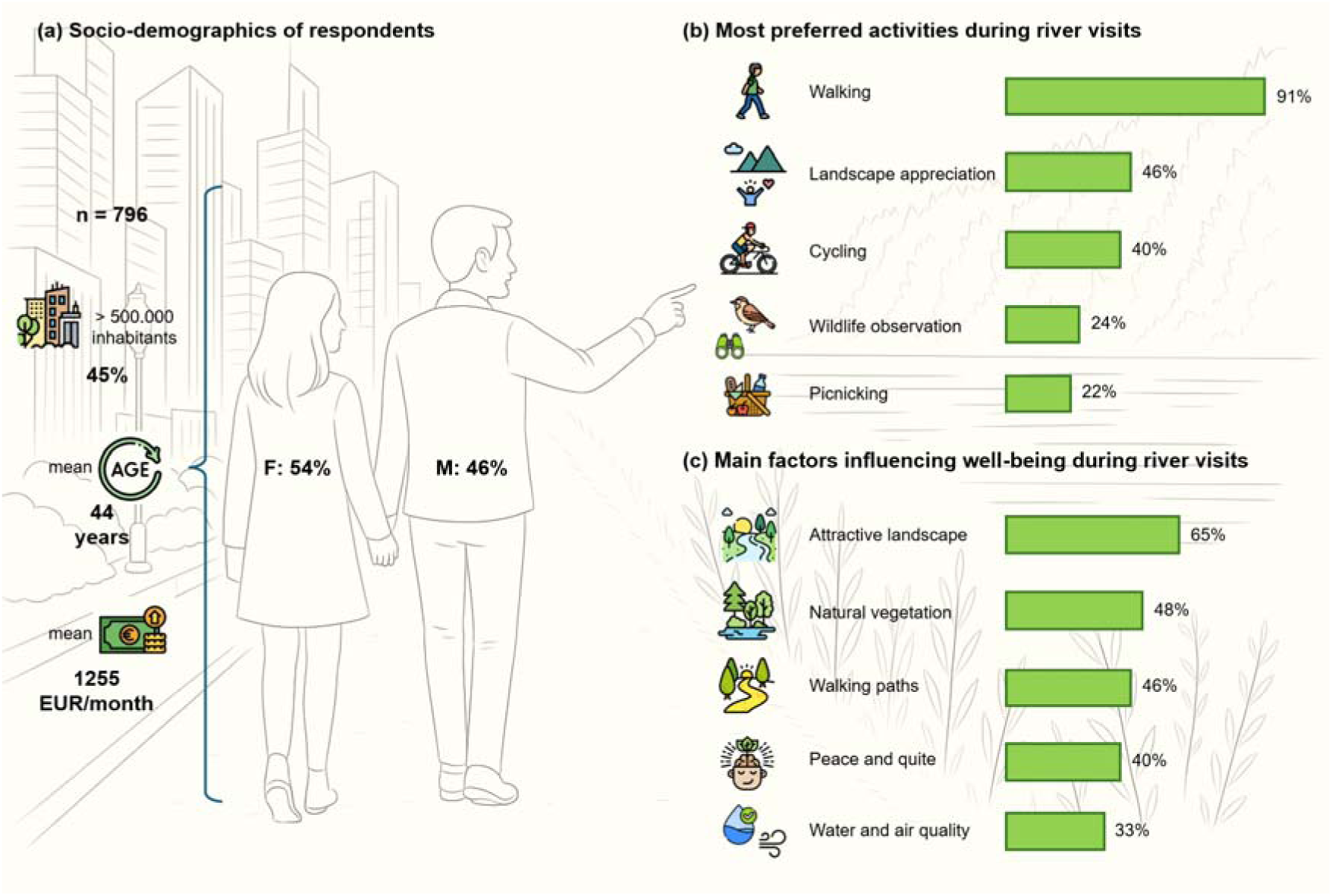
Summary of the main descriptive statistics. Panel (a) displays the main socio-demographics of the final sample (n=796). Most respondents lived in large cities, with 45% residing in municipalities of over 500,000 inhabitants. Just over half of respondents identified as women (54%), with an average age of 44 years throughout all respondents (SD = 13.5). Monthly net income per person averaged 1255 EUR/month, though nearly 10% of respondents chose not to disclose their income. Panel (b) displays respondents most preferred activities during river visits. Respondents reported visiting urban rivers an average of 19 times per year, though frequency varied widely (SD = 44.1). Panel (c) displays the respondents’ main factors influencing their well-being. Full descriptive statistics of the final sample are provided in the Supporting information (S2).

### 3.2 Willingness to pay for urban riparian attributes

Respondents were willing to pay for almost all attribute levels relative to the status quo, except for those associated with dead wood and the lowest level of management intensity (Figure 5).

**Figure 5:**
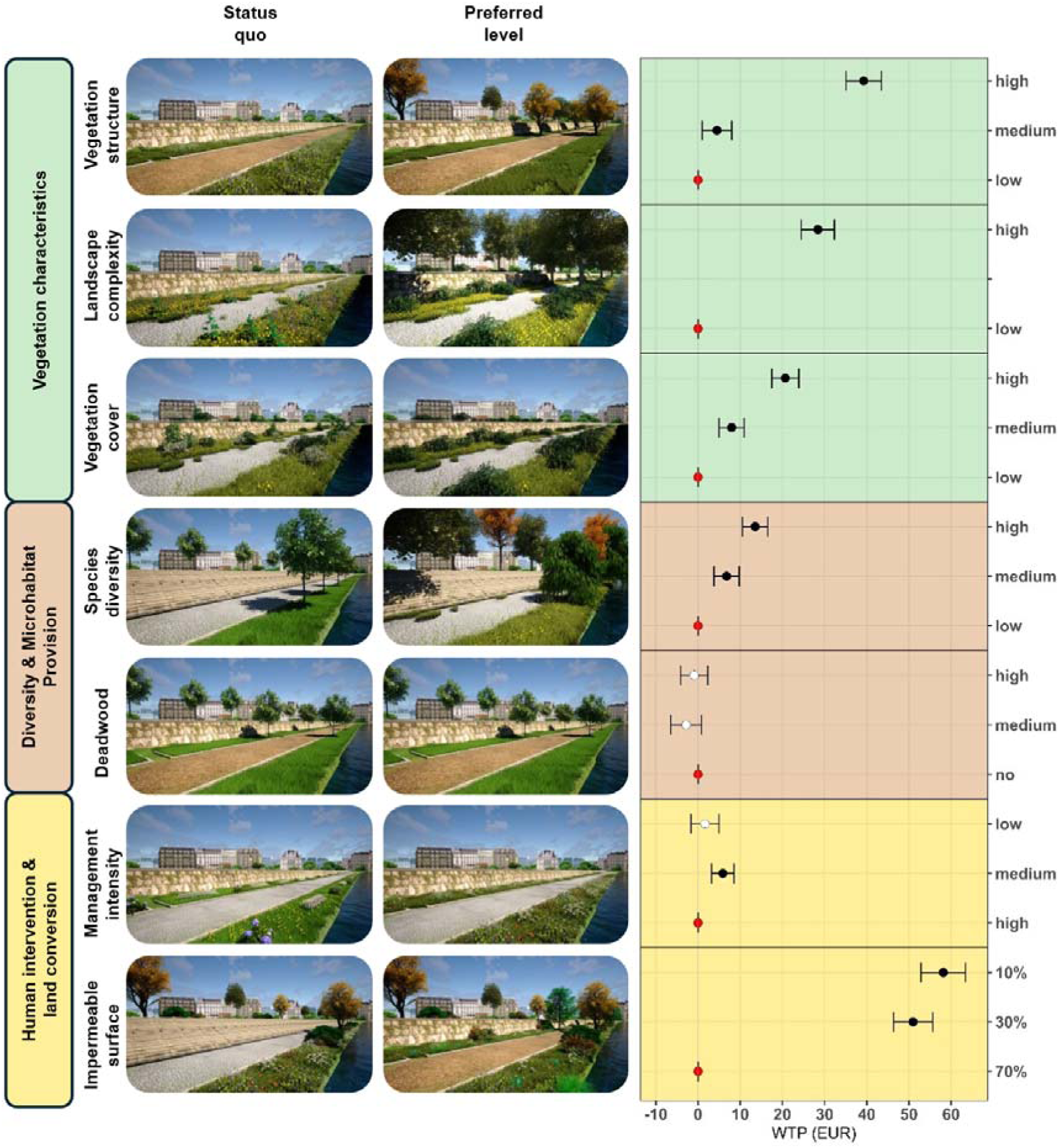
Pooled respondents’ willingness to pay (WTP) for management interventions in urban riparian areas. WTP (in EUR) and 95% confidence interval. Red dots indicate the reference level of each attribute, and black dots indicate significant differences from the reference level at the 95% confidence interval. The full results table is provided in the Supporting Information (S3).

Overall, respondents showed strong positive preferences for measures that enhance the naturalness, vegetation cover, and diversity of urban riparian areas. Among vegetation characteristics, the highest WTP was recorded for riparian areas with trees (€39.26), followed by intermediate structures with shrubs (€4.48), with low-structure vegetation (grass and herbaceous layer) serving as the reference level. A similar trend was observed for landscape complexity, with multiple vegetation layers strongly preferred (€28.41) over single-layer structures. For vegetation cover, higher density was consistently favored, with WTP of €20.70 for dense cover and €7.96 for moderate cover.

Preferences for species diversity followed a clear monotonic gradient. High species richness was associated with the highest WTP (€13.53), followed by medium richness (€6.77), with low species richness serving as the reference level. By contrast, the attribute dead wood showed no clear pattern. Although respondents associated increasing amounts of dead wood with negative WTP, with the strongest disapproval for the medium level (–€2.81), WTP for both levels did not differ statistically from the reference level (no deadwood).

Preferences regarding management intensity followed an inverted U-shape, with the highest WTP (€5.87) recorded for moderate-intensity regimes—defined as a mix of natural and planted vegetation maintained twice per year. The lack of a statistically significant difference in WTP for the lowest management intensity compared to the reference level suggests a high degree of polarization in preferences for this attribute. The difference in WTP for the lowest level of management intensity compared to the reference level was not statistically significant.

The most pronounced responses were observed for the attribute impermeable surface. Among all tested attributes, the highest WTP (58.15 EUR) was associated with the lowest share of concrete sealed area (10% sealed / 90% vegetated) along urban riparian areas, while even a moderate share (30% sealed / 70% vegetated) elicited substantial approval (50.97 EUR) compared to the reference level with a share of 70% sealed/ 30% vegetated. The full results are reported in the Supporting Information (S3).

### 3.3 Direct assessment of the preferences for the selected attributes and levels

To validate the preference study results, we conducted a direct preference ordering exercise in which respondents identified their most and least preferred levels for each attribute. This produced a clear ranking of attribute levels (Figure 6), closely matching the patterns observed in the DCE. In both approaches, respondents consistently favored more nature-based features. For example, 84.8% selected high-structure vegetation (trees, shrubs, grasses, and herbs) as most attractive, while 88.2% rated low-structure vegetation as least attractive. High species richness was preferred by 69–80% of respondents across vegetation contexts, whereas low richness was least attractive for 82–84%. Dense vegetation cover was favored by 65–69%, while low cover was least preferred by 83–86%. For landscape complexity, 82.8% preferred high complexity over low, and 84.2% preferred medium complexity over low.

**Figure 6:**
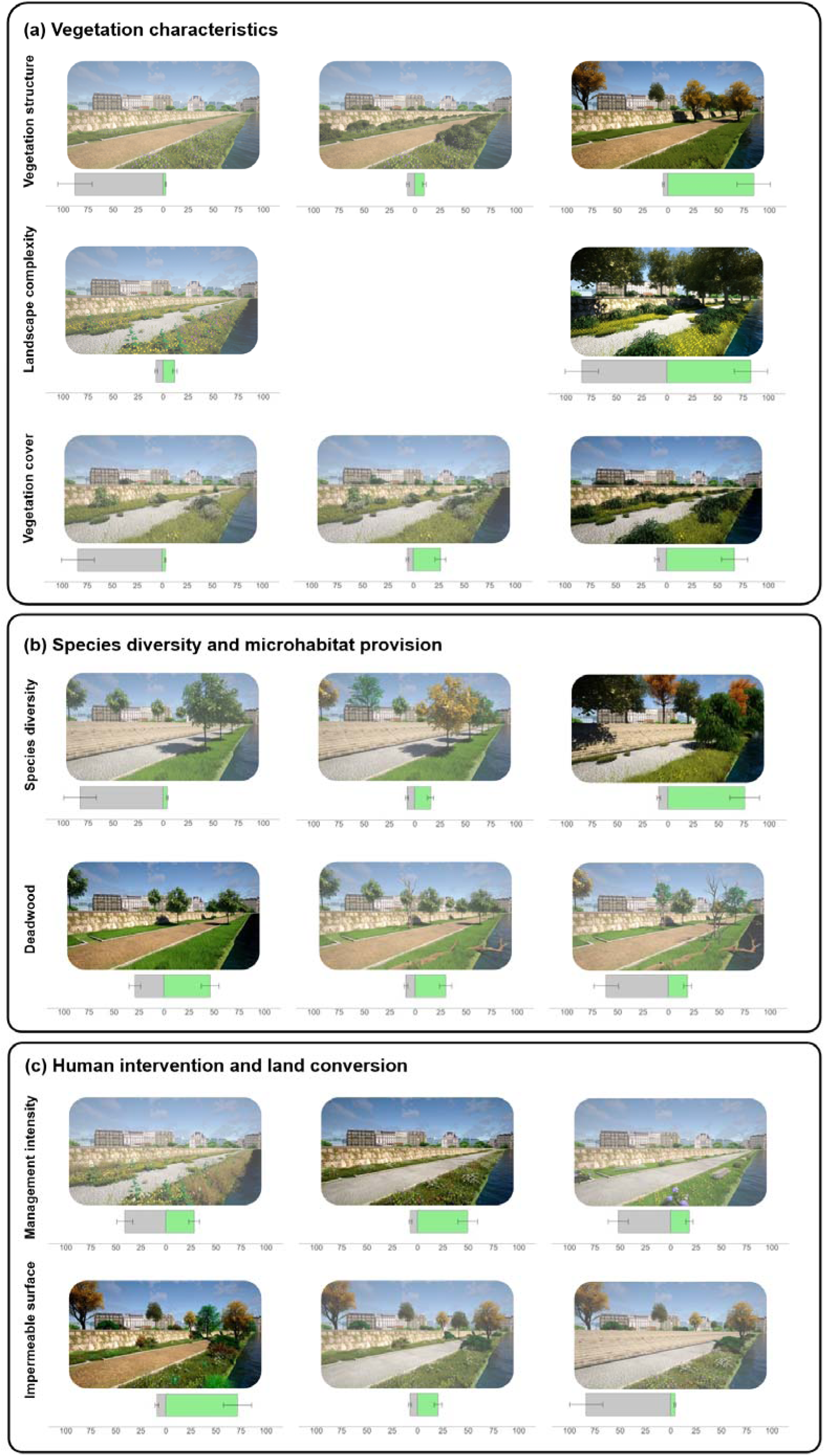
Validation of preferences for urban riparian characteristics. In green/gray the share of respondents who perceived a given level as the most/least attractive in percentages. Panel (a) shows all attributes within the category vegetation characteristics (vegetation type, landscape complexity and vegetation cover), panel (b) the shows all attributes within the category species diversity and microhabitat provision (species diversity and deadwood) and panel (c) shows all attributes within the category human intervention and land conversion (management intensity and impermeable surface). Example visualizations of each attribute level are given above the bar plots. Images with no transparency indicate the most preferred level of each attribute. The results table of the preference validation for each attribute is provided in the Supporting Information (S4).

Some attributes showed a higher degree of preference polarization. For dead wood, the absence of dead wood was most often preferred (46.1%), while medium levels were most often least preferred (61.4%), yet neither extreme attracted overwhelming consensus. These results align with the DCE findings, where mean WTP for medium and high levels did not differ statistically from the base level. This should not be interpreted as a lack of preference, but rather as evidence of strong polarization— yielding non-significant mean WTP but highly significant standard deviations in the normal distribution of WTP (see SI for details). A similar pattern emerged for management intensity, where moderate regimes—defined as a mix of natural and planted vegetation maintained twice per year—were most often preferred (49–50%), while high-intensity management was least preferred (47–56%), indicating divided opinions between low- and high-intensity regimes. The full results of the preference validation are provided in the Supporting Information (S4).

## Discussion

Urban riparian areas are multifunctional spaces where ecological processes and human use intersect, yet their planning and management often fail to account for the public’s specific preferences for different ecological features. While previous valuation studies have typically relied on broad proxies such as “naturalness,” our study advances this field by systematically decomposing biodiversity and its ecological function into a multidimensional set of observable attributes and quantifying their relative importance using a discrete choice experiment (DCE). This approach allows us to disentangle how the public values distinct aspects of riparian biodiversity — from vegetation composition and structural complexity to species richness and ground sealing — and to translate these preferences into monetary terms that can inform targeted, evidence-based urban river management.

We found strong and consistent preferences for attributes that signal high ecological quality: structurally complex vegetation, dense plant cover, high species diversity, and a substantial reduction in impermeable surfaces. These results were reinforced by a direct preference-ranking exercise, suggesting robust public support for biodiversity-oriented interventions in urban riparian areas.

### Vegetation structure and cover

Respondents expressed the highest WTP for tree-dominated riparian zones, followed by mixed shrub–tree assemblages, over low-structure herbaceous vegetation. Structural complexity and dense vegetation cover play critical roles in maintaining ecological function in riparian systems—providing habitat heterogeneity, shading to regulate stream temperatures, and connectivity between aquatic and terrestrial habitats (Groffman et al., 2003; Naiman et al., 2005; Peroni et al., 2023). Dense canopy and understory vegetation can also improve air quality, mitigate urban heat island effects, and buffer stormwater flows (Zhao et al., 2019; Munné et al., 2003).

The strong public support for these features suggests that restoring more natural vegetation structures is both ecologically desirable and socially acceptable. This aligns with a comparable study by Salm et al. (2023), who found that respondents preferred structural complex and diverse vegetation over monocultures, and additionally valued its ecological function in regulating urban temperatures. Further, our results align with earlier Polish evidence showing that urban residents are willing to pay for increased tree cover—such as street tree planting in Łódź (Giergiczny & Kronenberg, 2014)—and reinforce the potential for vegetation-based restoration to achieve both ecological and social goals in urban riparian management.

### Species diversity and microhabitat provision

Preferences followed a clear positive gradient for plant species richness, indicating that the public values interventions that enhance biodiversity directly. This is consistent with the ecological literature linking riparian diversity to ecosystem services such as bank stabilization, nutrient retention, and habitat provision (Naiman et al., 1993; Munro et al., 2009; Singh et al., 2021).

By contrast, dead wood elicited more mixed responses. While ecologists recognize its role in nutrient cycling, habitat provision, and overall ecosystem resilience (Maser & Trappe, 1984; Lassauce et al., 2011; Rimle et al., 2017), public preferences in our study did not indicate broad support for its presence in urban contexts. This contrasts with findings from non-urban forest settings, where respondents showed willingness to pay for ecological components like dead wood (Giergiczny et al., 2015). The divergence may reflect aesthetic expectations or perceived safety concerns in urban environments (Nassauer, 1995; Qiu et al., 2013), suggesting that ecological features valued in wilderness contexts are not automatically transferable to urban settings. Public education and interpretation could help bridge this gap, enhancing awareness of the ecological benefits of dead wood in urban riparian areas.

### Human intervention and land conversion

Respondents preferred moderately managed riparian landscapes over both high- and low-intensity regimes, suggesting a desire for a balance between ecological function and a degree of visual order. This aligns with research showing that perceptions of “care” influence public acceptance of urban biodiversity interventions (Nassauer, 1995; Qiu et al., 2013).

Most strikingly, the highest WTP across all attributes was for reducing impermeable surfaces. This preference resonates with concerns about the ecological impacts of soil sealing—altered hydrological regimes, degraded water quality, habitat loss, and exacerbated flood risks (Walsh et al., 2005; Meyer et al., 2005; Serra et al., 2019). Impervious surfaces also contribute to urban heat islands and disrupt groundwater recharge (Fini et al., 2017; Sousa et al., 2024). Our results are, to our knowledge, the first DCE evidence directly quantifying public preferences for impermeable surface reduction in urban riparian areas. They align with findings by Johnson & Geisendorf (2022), who observed strong public valuation of water quality improvements closely tied to reduced soil sealing.

### Implications and limitations

Our findings strengthen the case for integrating biodiversity conservation explicitly into urban riparian planning, supported by public willingness to finance interventions. The evidence for preferences favoring structurally complex vegetation, high species richness, and reduced impermeable surfaces provides actionable guidance for urban planners aiming to reconcile ecological goals with social acceptance.

Nonetheless, stated preference methods like DCEs face well-known limitations, including hypothetical and framing biases, information overload, and social desirability effects (Hensher, 2010; Rolfe et al., 2002; Lopez-Becerra & Alcon, 2021; Shr et al., 2019). We mitigated these risks by providing clear attribute descriptions, realistic visualizations, and sufficient reflection time. By employing multiple biodiversity indicators rather than a single proxy concept, we also addressed the conceptual complexity of biodiversity in economic valuation (Bartkowski et al., 2015; Díaz et al., 2018; Hill et al., 2021; Kenter et al., 2015). Still, behavioral validity cannot be fully guaranteed (Sakurai & Uehara, 2023).

Urban rivers in Poland—like many in Europe—reflect centuries of human modification for flood control, navigation, and urban development (Brown et al., 2018; Kubiak-Wójcicka et al., 2017). Yet, restoration efforts, including re-naturalization and blue– green infrastructure expansion, are increasingly recognized for their ecological and social value (Gilvear et al., 2013; Dai et al., 2021). Unfortunately, the ways in which evidence drawn from valuation studies is incorporated into current urban planning still lags behind. Moreover, planning decisions rarely account for the diversity of public preferences, causing potential conflicts over use, safety or aesthetics.

Thus, we argue that the success of interventions in urban riparian areas depends strongly on the integration of both monetary and non-monetary values (ecological, social, cultural) to inform urban planning and changing the paradigm so that natural features are not just ornaments but core features that shape how cities grow and function. Embedding biodiversity concerns into urban river planning, while ensuring local stakeholder and diverse user groups engagement (Quintero-Uribe et al., 2022; Dunn-Capper et al., 2023), will be essential to achieve climate-resilient, socially equitable, and ecologically functional urban riparian systems.

## Supporting information

Supporting_Information_1_Questionnaire_english

## Author contributions

Arpad Erik Thoma and Marek Giergiczny conceived the idea for the research; Arpad Erik Thoma and Marek Giergiczny designed the methodology, Marek Giergiczny led the data analysis; Arpad Erik Thoma and Marek Giergiczny wrote the manuscript with substantial input from Henrique M. Pereira.

## Acknowledgements

Arpad Erik Thoma would like to thank the working group Biodiversity Conservation at the German Centre for Integrative Biodiversity Research (iDiv) for their support. Marek Giergiczny acknowledges the financial support from the National Science Centre in Poland (2021/43/B/HS4/03371) and the iDiv Flexpool Funds.

## Data availability statement

The data used in this study were collected from online survey responses; Anonymised choice experiment data and results may be found in Zenodo: https://zenodo.org/records/16882192

## Supporting Information

Additional supporting information can be found online in the Supporting Information section at the end of this article.

Supporting Information **S1**: Questionnaire

Supporting Information **S2**: Descriptive statistics of respondents in Poland (N = 796)

Supporting Information **S3**: Respondents preference results

Supporting Information **S4**: Validation of attribute preferences

