## Supporting_Information_2-4 for "Greener, Wilder, Better: Valuing Complexity and Diversity in Urban Riparian Areas"

**This document contains the following Supporting Information for this article:**

Supporting Information **S2**: Descriptive statistics of respondents in Poland (N = 796)

Supporting Information **S3**: Respondents preference results

Supporting Information **S4**: Validation of attribute preferences

Supporting Information **S2**: Descriptive statistics of respondents in Poland (N = 796)

| | Poland | | | |
|---|---|---|---|---|
| Respondent description | Mean | SD | Min | Max |
| Age | 43.7 | 13.5 | 18 | 75 |
| Gender (Women) | 0.54 | - | - | - |
| Income (Monthly EUR) | 1253.57 | 668.57 | 476.19 | 4761.90 |
| Respondents not reporting income | 0.094 | - | - | - |
| Higher education | 0.43 | - | - | - |
| Number of river visits per year | 18.6 | 44.1 | 0 | 365 |

Supporting Information **S3**: Respondents preference results

Results of the mixed logit model for Poland (N = 796). Estimates for the mean, standard deviation (SD), and standard errors (SE). Results significant at the 5% level are in bold.

| Level | Mean | SE | SD | SE |
|---|---|---|---|---|
| SQ | **-6.45284** | 0.404114 | **12.99987** | 0.758015 |
| Medium - Vegetation type - shrubs | **0.381692** | 0.151552 | **0.70143** | 0.219656 |
| High - Vegetation type - trees | **3.341693** | 0.183322 | **3.026092** | 0.173728 |
| High - Landscape complexity | **2.41772** | 0.169649 | **2.337493** | 0.189901 |
| Medium - Vegetation cover | **0.677341** | 0.127609 | **0.301275** | 0.169273 |
| High - Vegetation cover | **1.761477** | 0.139339 | **1.118984** | 0.142941 |
| Medium – Species Diversity | **0.576242** | 0.129082 | 0.368303 | 0.246166 |
| High – Species Diversity | **1.151771** | 0.130714 | **0.617315** | 0.187566 |
| Medium – Dead wood | -0.23955 | 0.1586 | **0.898892** | 0.240041 |
| High – Dead wood | -0.07597 | 0.141039 | 0.04714 | 0.225325 |
| Medium - Management intensity | **0.499819** | 0.115478 | 0.327393 | 0.204652 |
| Low - Management intensity | 0.140487 | 0.142465 | **2.394719** | 0.170843 |
| 30% impermeable surface | **4.337736** | 0.201506 | 0.343241 | 0.252951 |
| 10% impermeable surface | **4.948722** | 0.23001 | **3.138627** | 0.177311 |
| Tax (cost) | **-0.91623** | 0.049379 | **0.663047** | 0.042784 |

**Model diagnostics** LL at convergence -9011,13
LL at constant(s) only -13790,7
McFadden's pseudo-R² 0,346577
Ben-Akiva-Lerman's pseudo-R² 0,51598
AIC/n 1,419776
BIC/n 1,43733
n (observations) 12736
r (respondents) 796
k (parameters) 30

Supporting Information **S4**: Validation of attribute preferences

Results of the direct attribute preferences (N = 796). Most and least attractive attribute levels are indicated in percentages, with the most or least attractive level in bold.

| Attribute | level | most attractive | least attractive |
|---|---|---|---|
| Vegetation structure | low | 2.49 | **88.17** |
| | medium | 9.27 | 7.14 |
| | high | **84.75** | 4.68 |
| | not rel. | 3.5 | NA |
| Landscape complexity | low | 11.53 | 7.06 |
| | high | **82.82** | **84.24** |
| | not rel. | 5.65 | 8.71 |
| Vegetation cover in medium vegetation structure | low | 3.05 | **83.02** |
| | medium | 28.25 | 5.42 |
| | high | **64.52** | 11.56 |
| | not rel. | 4.18 | NA |
| Vegetation cover in high vegetation structure | low | 3.28 | **85.83** |
| | medium | 24.97 | 6.62 |
| | high | **69.04** | 7.55 |
| | not rel. | 2.71 | NA |
| Species diversity in low vegetation structure | low | 4.42 | **83.81** |
| | medium | 19.47 | 7.62 |
| | high | **69.03** | 8.57 |
| | not rel. | 7.08 | NA |
| Species diversity in medium vegetation structure | low | 3.54 | **83.33** |
| | medium | 11.5 | 5.56 |
| | high | **80.53** | 11.11 |
| | not rel. | 4.42 | NA |
| Species diversity in high vegetation structure | low | 4.42 | **81.98** |
| | medium | 15.93 | 9.91 |
| | high | **77.88** | 8.11 |
| | not rel. | 1.77 | NA |
| Dead wood | none | **46.1** | 29.17 |
| | low | 29.94 | 9.4 |
| | medium | 18.87 | **61.43** |
| | not rel. | 5.08 | NA |

| | | | | |
|---|---|---|---|---|
| Management intensity in low vegetation structure | low | 30.62 | | 37.43 |
| | medium | **49.94** | | 6.97 |
| | high | 15.41 | | **55.61** |
| | not rel. | 4.29 | NA | |
| Management intensity in medium vegetation structure | low | 28.7 | | 39.5 |
| | medium | **49.6** | | 7.55 |
| | high | 17.51 | | **52.95** |
| | not rel. | 4.18 | NA | |
| Management intensity in high vegetation structure | low | 25.54 | | 46.18 |
| | medium | **48.59** | | 6.93 |
| | high | 22.03 | | **46.89** |
| | not rel. | 3.84 | NA | |
| Impermeable surfaces | low | **71.53** | | 8.83 |
| | medium | 20.23 | | 7.54 |
| | high | 4.18 | | **83.63** |
| | not rel. | 4.07 | NA | |
