## Supporting_Information_1_Questionnaire_english for "Greener, Wilder, Better: Valuing Complexity and Diversity in Urban Riparian Areas"

**This document contains the following Supporting Information for this article:**

Supporting Information **S1**: English version of the questionnaire

We would like to invite you to participate in a research study conducted by the University of Warsaw. Our aim is to investigate perceptions of riverside vegetation in Polish cities located along rivers.

No specialist knowledge is required to participate in the survey. There are no right or wrong answers in the survey. We are interested in all the opinions expressed.

Participation in the study involves carefully reading the information contained in the survey and takes about **20 minutes**.

**This is a scientific study, the results of which can be used to change the way urban greenery is managed, including in the city where you live.**

Therefore, if you are unable to devote 20 minutes to carefully reading the survey, please do not participate.

The survey is anonymous and the data collected will only be analysed at an aggregate level.

#### Metrics

**M1. What is your year of birth?** (*One answer.*)

**M2. Please indicate your gender.** (*One answer.*)
Options:
1) Male,
2) Female,
3) Other

**M3. What is your employment status?** (*One answer*)
Options:
1) Full-time job;
2) Part-time job;
3) Trainee;
4) Unemployed;
5) Self-employed;
6) Retired;
7) Not working, studying
8) Other what?

**M4. What is your personal monthly net income (in PLN)?** (*One answer.*)
Options:
1) Less than 2,000;
2) 2 001-3 000;
3) 3 001-4 000;
4) 4 001-5 000;
5) 5 001-6 000;
6) 6 001-7 000;
7) 7 001-8 500;
8) 8 501-10 000;
9) 10 001-15 000;
10) 15 001-20 000;
11) more than 20 000;
12) Refusal to answer.

**M5. What is your marital status?** (*One answer.)*
Options:
1) Married;
2) Divorced
3) Living in a civil partnership
4) Single
5) Separated.

**M6. What is your educational background?** (*One answer.*)
Options:
1) Primary or lower secondary school,
2) Basic vocational,
3) Secondary,
4) Post-secondary,
5) Higher bachelor's degree
6) Master's degree

**M7. What is the size of the locality in which you live?** (*One answer.*)
Options:
1) City with less than 20,000 inhabitants
2) City with 20 001- 100 000 inhabitants
3) City with 100 001- 500 000 inhabitants
4) City with more than 500,000 inhabitants

**M8. What is the postcode of your place of residence?**
____________ (*Please fill in*)

**//END OF PAGE//**

**SECTION B -** ***Perception and recreation***

In this section we would like to ask you about your perception of the riverside area where you live.

**B1. Do you live in a locality with a river(s)?** (expandable)
Options: Yes; No

If NO, discontinue the interview.

**B2. Which of the following landscapes best represents the riparian zone where you live?** (random order)

Options: A; B; C; D; None

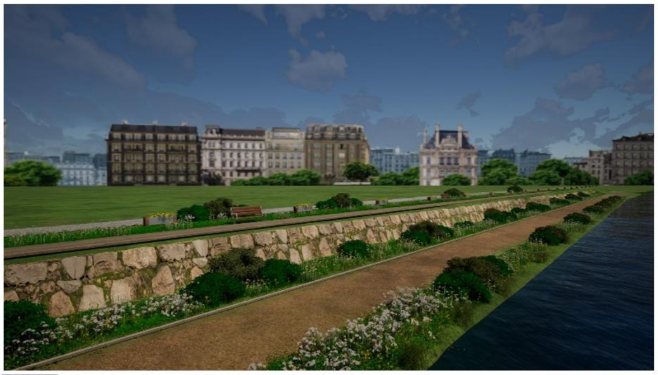

A

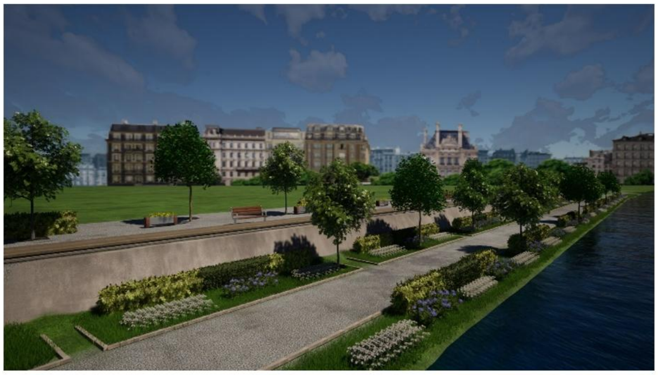

B

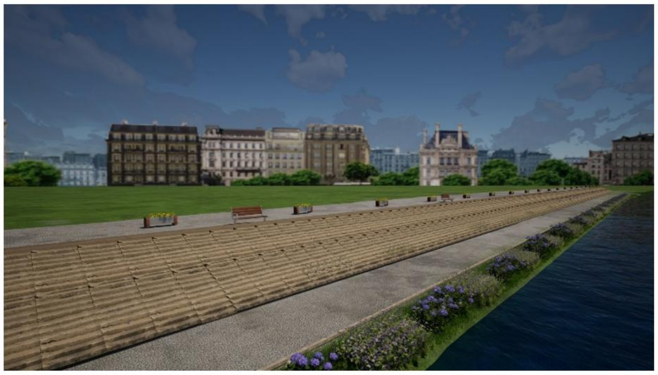

C

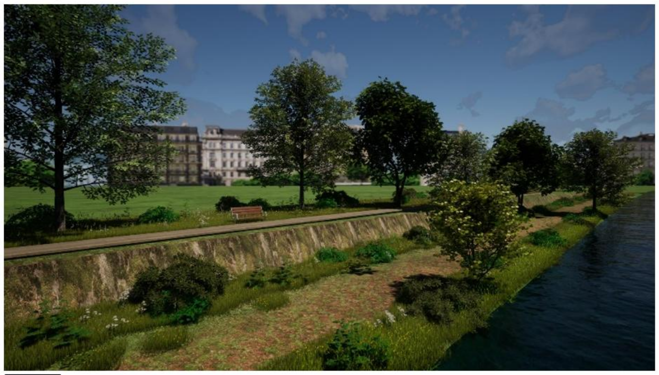

D

**B3. Please rank the following landscapes from most (1) to least attractive (4).** (random order)

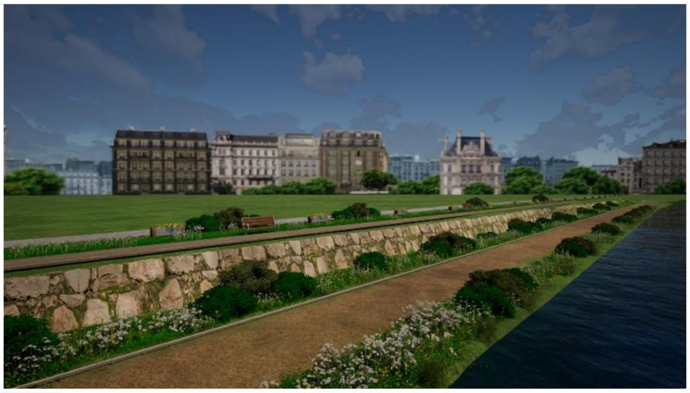

A

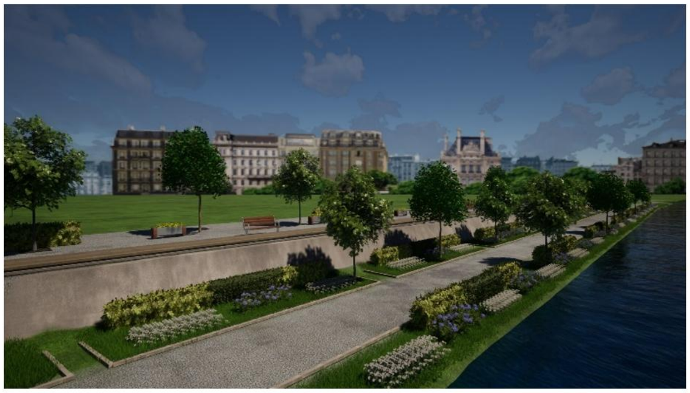

B

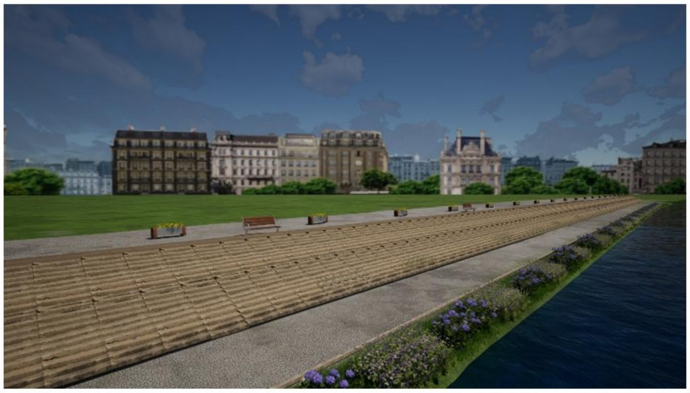

C

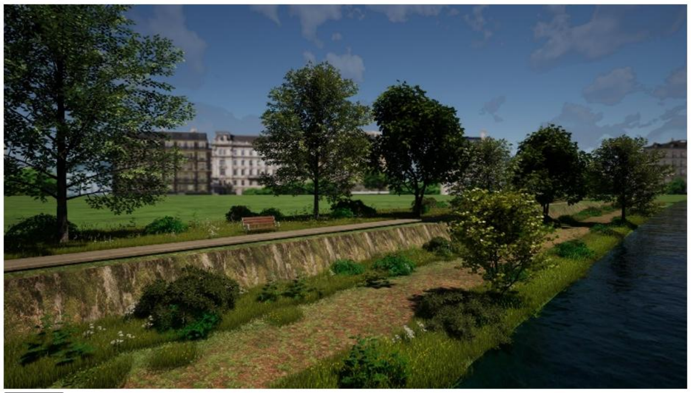

D

**B4. Approximately how far is the nearest riverside area from your home?**

a) Closer than 0.5 km
b) 0.5-1 km
c) 1-2 km
d) 2-3 km
e) Further than 3 km
f) Don't know, hard to say

**B4. In the last 12 months, have you visited the river area in your place of residence for recreational purposes?**
Options: Yes; No

B5-B7 only if in B4 answer. **Yes**

**B5. Approximately how many times have you visited the riverside area in the last 12 months?**
_________ times

**B6. What did you do during these visits to the river? Please tick all matching answers.** *[multiple answers].*

a) Walking
b) Running
c) Cycling
d) Observing the landscape
e) Fishing
f) Bird/wildlife watching
g) Picnic
h) Swimming/swimming
i) Sunbathing
j) Kayaking
k) Boating
l) Inne:_______________

**B7. Which of the following make you feel good about visiting the river. In your opinion, please select the three most important and rank them from most to least important to you** (please select and rank)

- Attractive landscape
- Water and air quality
- Presence of walking paths
- Presence of cycle paths
- Presence of recreational and leisure facilities (benches, shelters, exercise areas)
- Natural vegetation
- Presence of birds and other wildlife
- Peace and quiet
- Presence of restaurants, bars, cafes
- Inne:_________________

**//END OF PAGE//**

**INTRODUCTION**
**(ALL RESPONDENTS)**

Along the river there is a transition zone between land and water, referred to as the riparian zone. Due to the large differences in moisture content, natural riparian areas are covered with a variety of lush vegetation.

Riparian areas can vary in the way they are developed. Below are examples from various European cities.

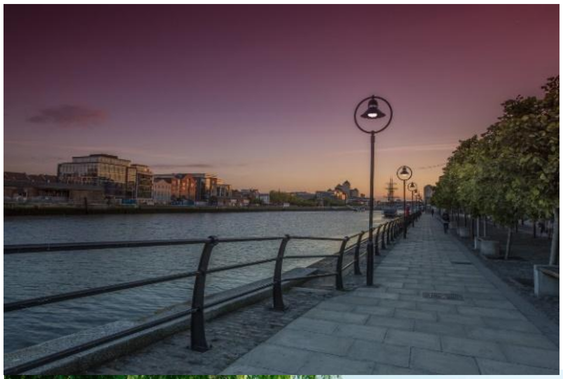
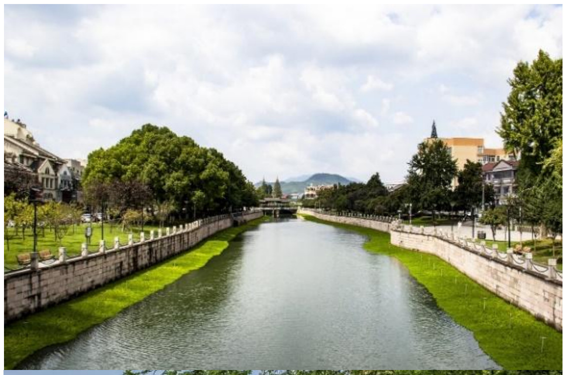
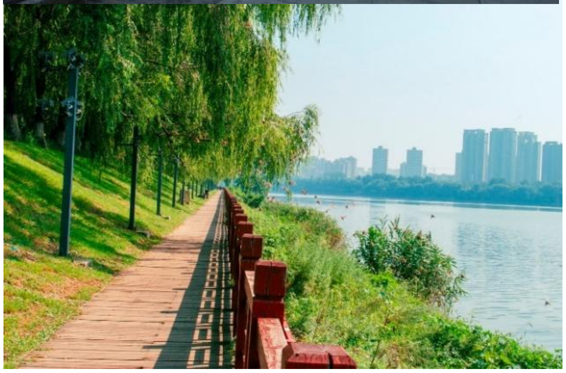
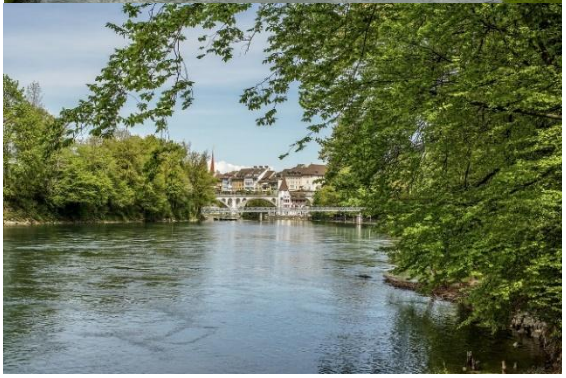

**//END OF PAGE//**

### Riparian vegetation management

In many cities, the riverside zone and its associated natural vegetation is undergoing significant transformation. The most important of these are:

Replacement of natural riverside vegetation by manicured ornamental vegetation.

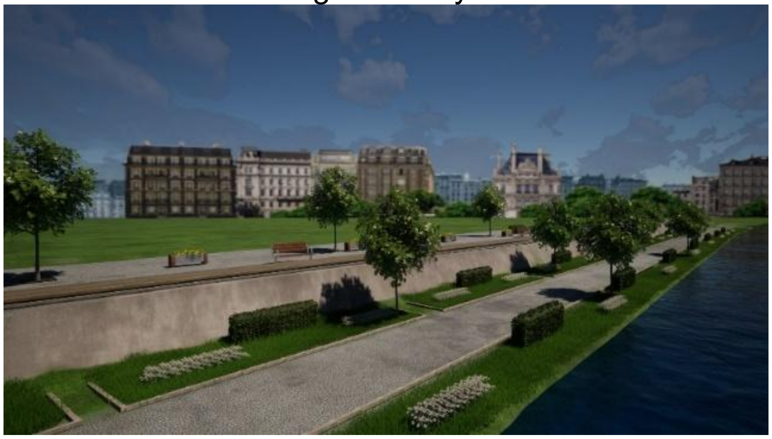

Replacement of natural surfaces (soil, gravel, sand) with hardened impermeable surfaces such as concrete or asphalt.

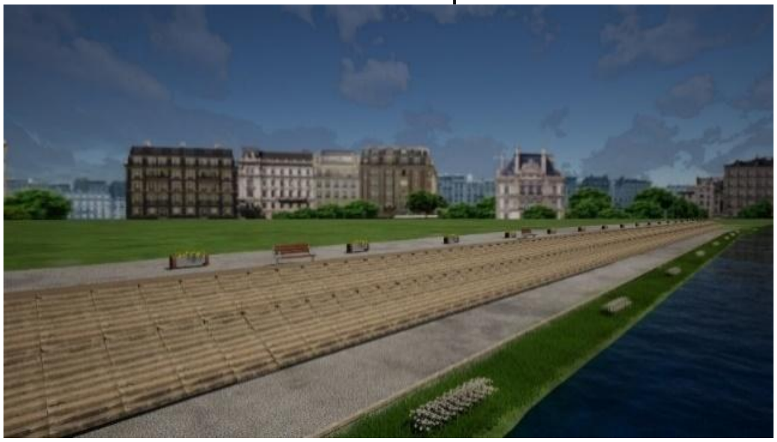

Removal from the riverside landscape of trees that have, for example, withered or been knocked down by the wind, for natural decomposition.

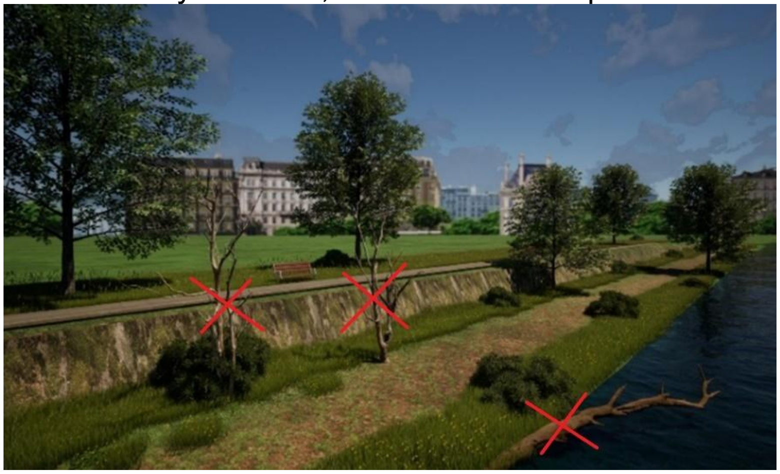

-

The main aim of our survey is to gauge opinions as to what type of vegetation and development of the riverside zone would be most attractive to the inhabitants of towns and cities along large rivers in Poland.

In order to assess the different management methods, we have created visualisations/photographs showing the appearance of the riverside zone for different vegetation types and management intensities.

**The study will consist of two parts.**
**The riverside landscape was described using the following characteristics:**

- **Vegetation type**
- **Vegetation density**
- **Species diversity**
- **Diversity of vegetation types**
- **Presence of impermeable surfaces** (e.g. concrete, asphalt)
- **Presence of dead wood**
- **Development intensity**

Each of these features will be described in detail in a moment.

**//END OF PAGE//**

We will now discuss each of the characteristics studied.

**//END OF INTRODUCTION//**

**SECTION C -** ***Perception and management of urban riparian vegetation***

#### VEGETATION TYPE

To describe the vegetation in the urban riparian zone, we assumed that different vegetation types correspond to different heights. We defined three levels:

- **Low vegetation** (e.g. grasses and herbs)
- **Medium vegetation** (shrubs)
- **High vegetation** (trees)

Examples of riparian areas differing in vegetation types:

| | | |
|---|---|---|
| 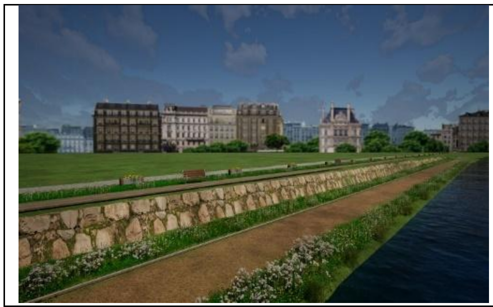 | 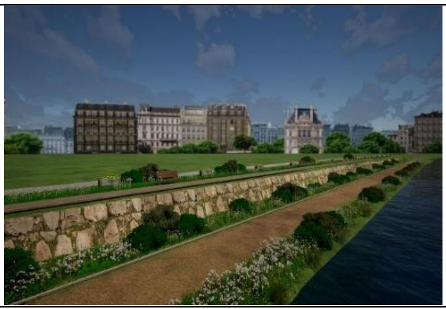 | 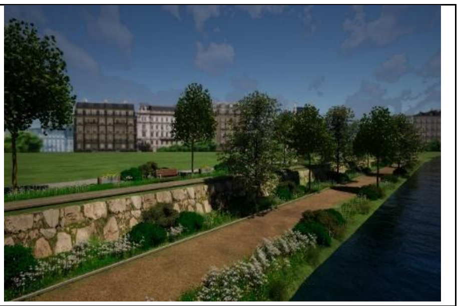 |
| Low<br>(e.g. grasses and herbs) | Medium<br>(shrubs, grasses and herbs) | High<br>(trees, shrubs, grasses and herbs) |

Higher vegetation types are accompanied by lower vegetation types, i.e. medium vegetation is accompanied by low vegetation and high vegetation is accompanied by both medium and low vegetation.

**C1. Which of the landscapes shown do you find the most attractive?**
**C2. Which of the landscapes shown do you find least attractive?**

**☐ Differences in vegetation type do not matter to me**

#### SPECIES DIVERSITY

The next characteristic we will consider is species diversity. It describes the number of species present in each vegetation type.

Each vegetation type (i.e. herbs and grasses / shrubs / trees) in our study can have three levels of species diversity: **low, medium** and **high.**

These levels correspond to the following landscape types:

**Species diversity**
(Example for low vegetation)

| | |
|---|---|
| 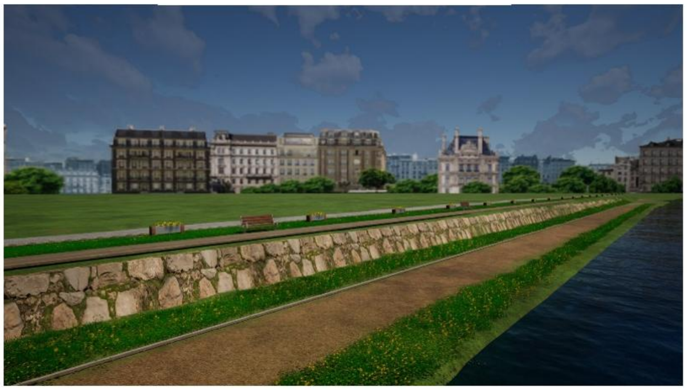 | **Low number of species**, vegetation type has low diversity |
| 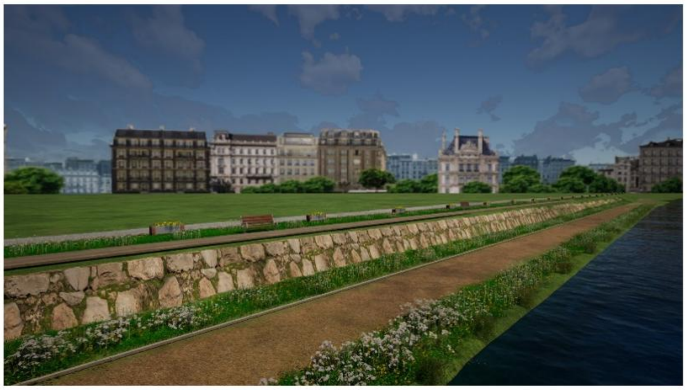 | **Medium number of species**, vegetation type is moderately diverse |
| 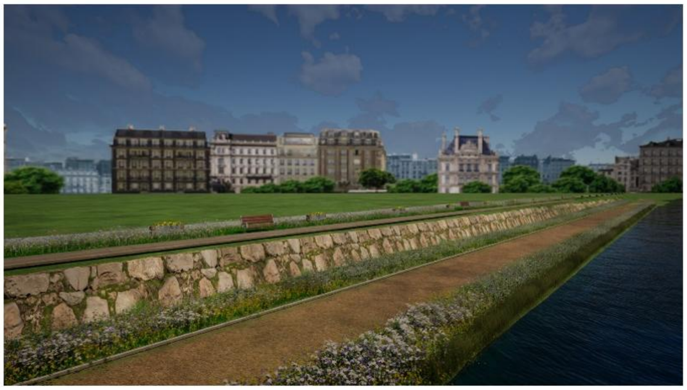 | **High number of species**, vegetation type is significantly differentiated |

**C3. Which of the landscapes shown do you find most attractive?**
**C4. Which of the landscapes shown do you find least attractive?**

☐ **The differences in species diversity do not matter to me.**

**Species diversity**
(Example for medium vegetation)

| | |
|---|---|
| 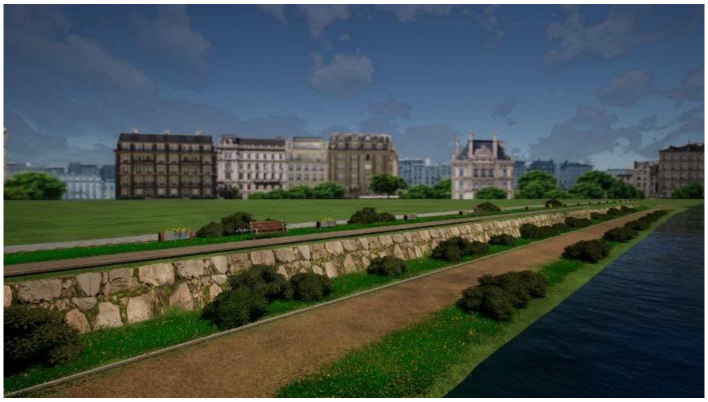 | **Low number of species**, vegetation type has little diversity |
| 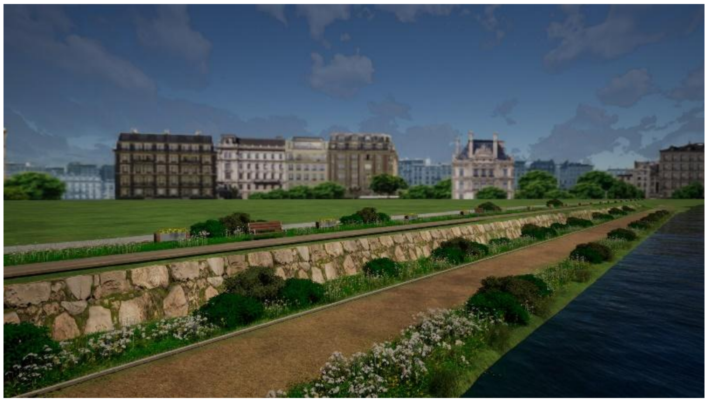 | **Medium number of species**, vegetation type is moderately diverse |
| 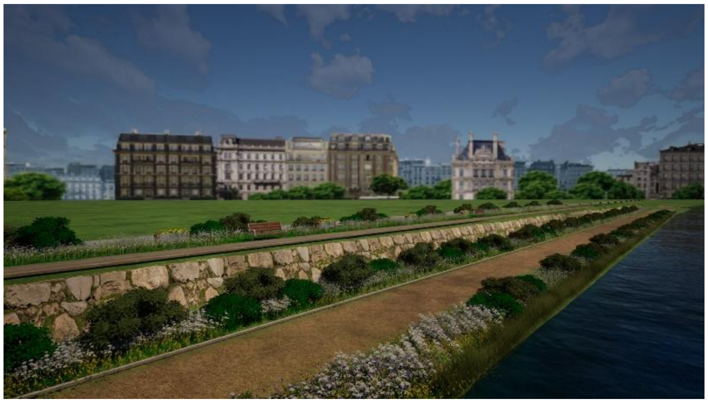 | **High number of species**, vegetation type is significantly differentiated |

**C3. Which of the landscapes shown do you find most attractive?**
**C4. Which of the landscapes shown do you find least attractive?**

☐ **The differences in species diversity do not matter to me.**

**Species diversity**
(Example for tall vegetation)

| | |
|---|---|
| 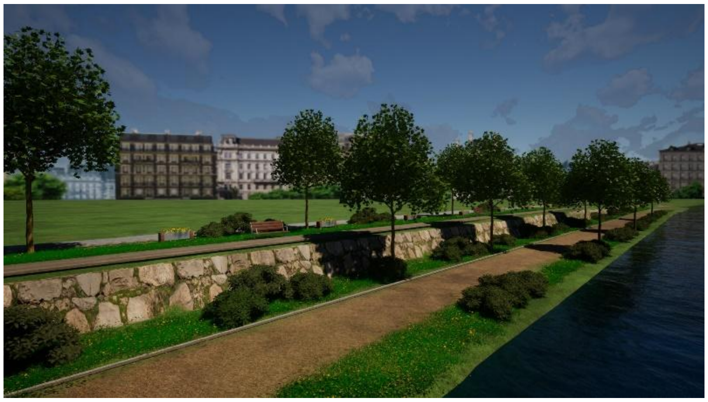 | **Low number of species**, vegetation type has little diversity |
| 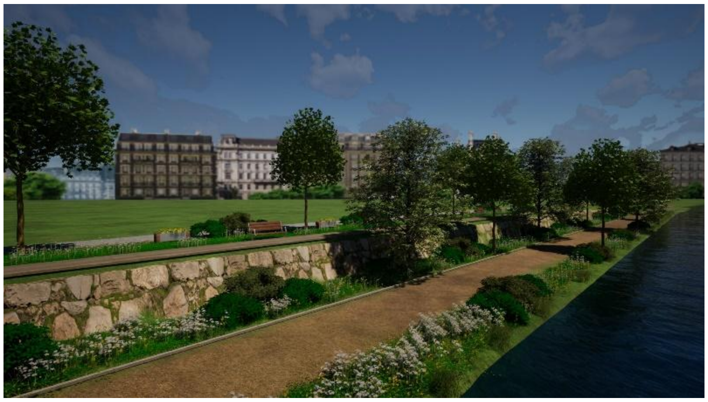 | **Medium number of species**, vegetation type is moderately diverse |
| 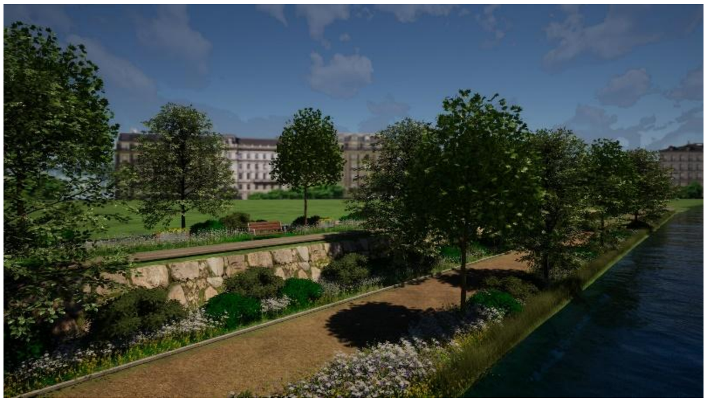 | **High number of species**, vegetation type is significantly differentiated |

**C3. Which of the landscapes shown do you find most attractive?**
**C4. Which of the landscapes shown do you find least attractive?**

**☐ The differences in species diversity do not matter to me.**

**VEGETATION COVER**

The next feature we will consider is vegetation density. We have distinguished three levels of vegetation density: low, medium and high density.

These levels correspond to the following landscape types:

**Vegetation cover**
(Example for shrubs)

| | |
|---|---|
| 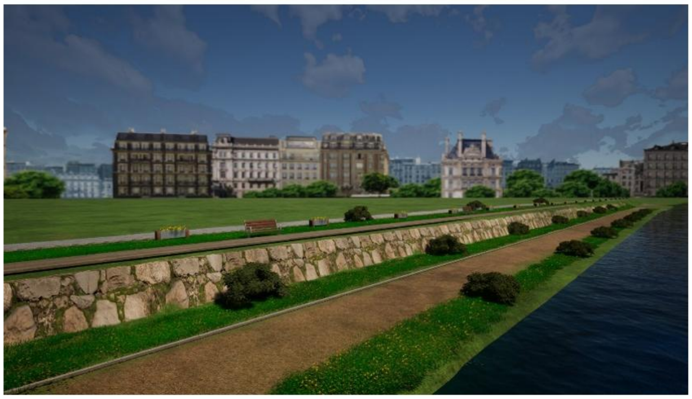 | **Low density**<br>**Low** vegetation cover |
| 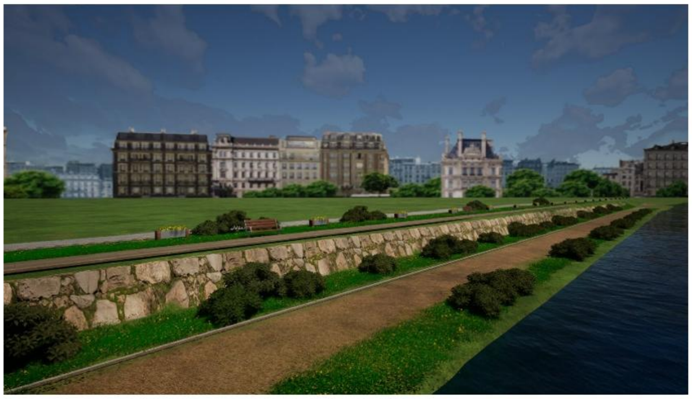 | **Medium density**<br>**Medium density Medium** vegetation cover |
| 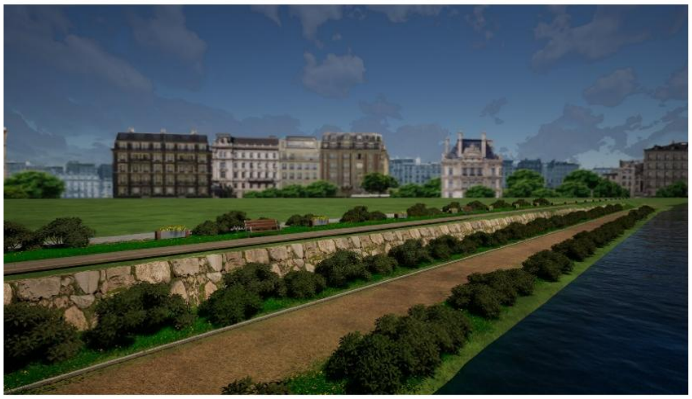 | **High density**<br>**High** vegetation cover |

**C5. Which of the landscapes shown do you find most attractive?**
**C6. Which of the landscapes shown do you find least attractive?**

**☐ Differences in vegetation density do not matter to me.**

**Vegetation cover**

Example for shrubs and trees:

| | |
|---|---|
| 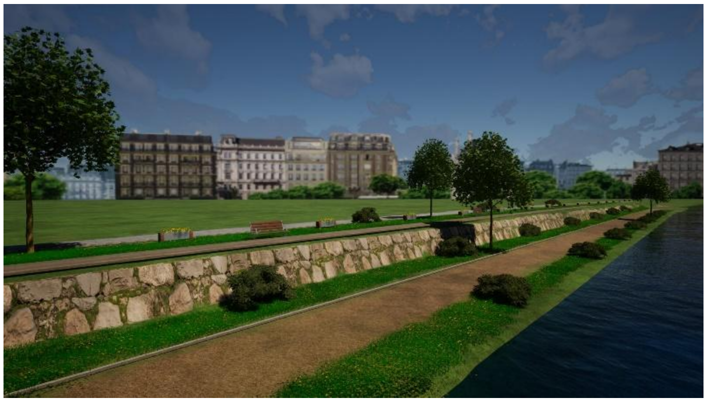 | **Low density**<br>**Low** vegetation cover |
| 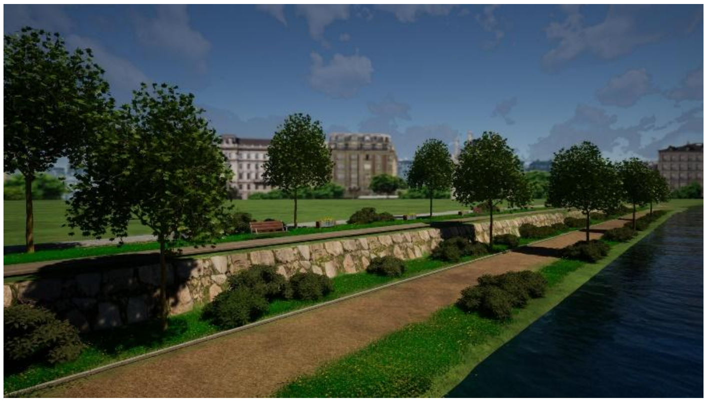 | **Medium density**<br>**Medium density Medium**<br>vegetation cover |
| 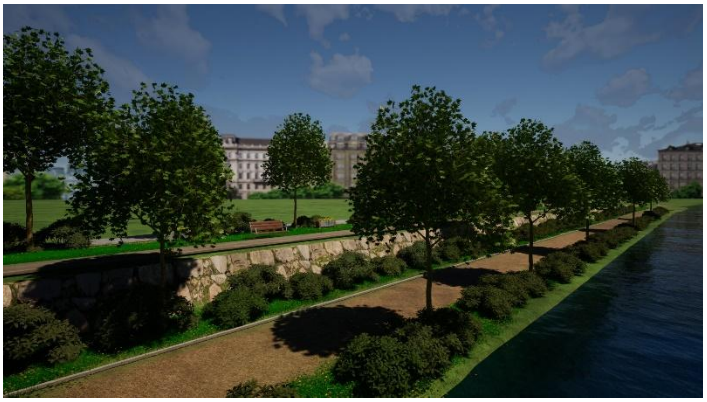 | **High density**<br>**High** vegetation cover |

**C5. Which of the landscapes shown do you find most attractive?**
**C6. Which of the landscapes shown do you find least attractive?**

☐ **Differences in vegetation density do not matter to me.**

**LANDSCAPE COMPLEXITY**

The next feature we will consider is the diversity of vegetation types. We have distinguished two levels of diversity: low and high.

Example for trees

<table>
<tr><td>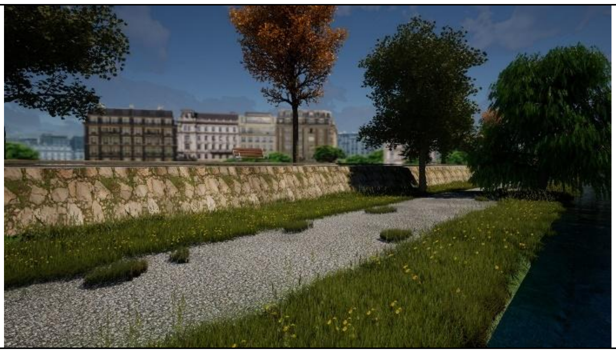</td><td>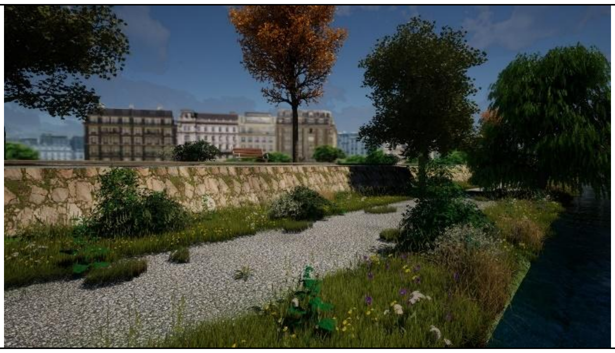</td></tr>
<tr><td>Low complexity: no shrub layer, poor low vegetation layer</td><td>High complexity: all vegetation types present (trees, shrubs and diverse low vegetation)</td></tr>
</table>

**Which of the landscapes shown do you find more attractive?**

**☐ Differences in vegetation density do not matter to me.**

Example for shrubs

<table>
<tr><td>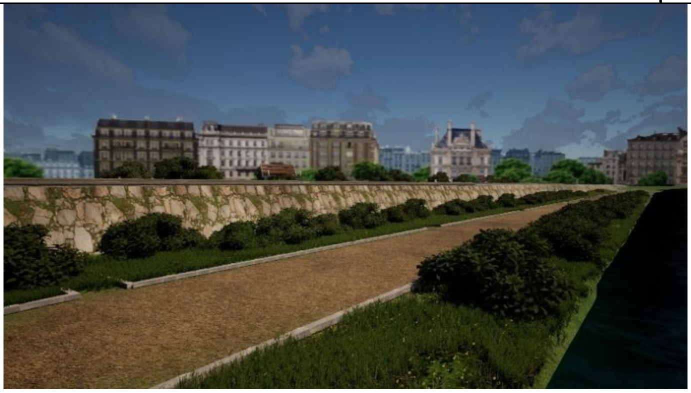</td><td>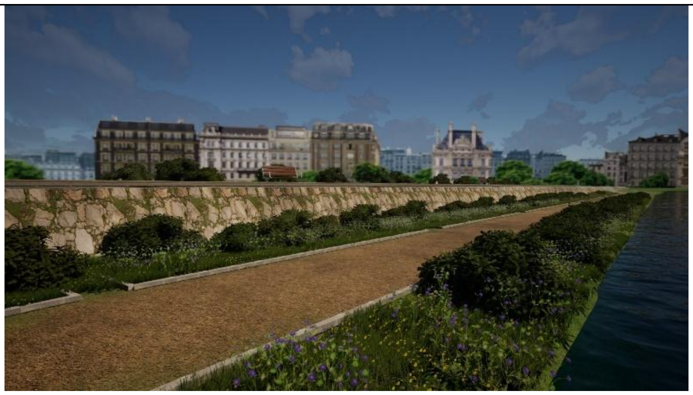</td></tr>
<tr><td>Low complexity: shrubs are accompanied by a poor understorey of low vegetation (no species diversity among grasses and herbaceous vegetation)</td><td>Medium complexity: shrubs are accompanied by a diverse understorey of low vegetation (high species diversity among grasses and herbaceous vegetation)</td></tr>
</table>

#### WOOD LEFT TO DECOMPOSE NATURALLY

Please now read the following information on dead wood.

Deadwood describes the amount of withered trees or dead wood that is left in the landscape to decompose naturally.

Currently, withered and dead trees are removed from riparian areas, mainly for aesthetic reasons. One of the aims of this survey is to investigate how such trees are perceived in the riparian landscape.

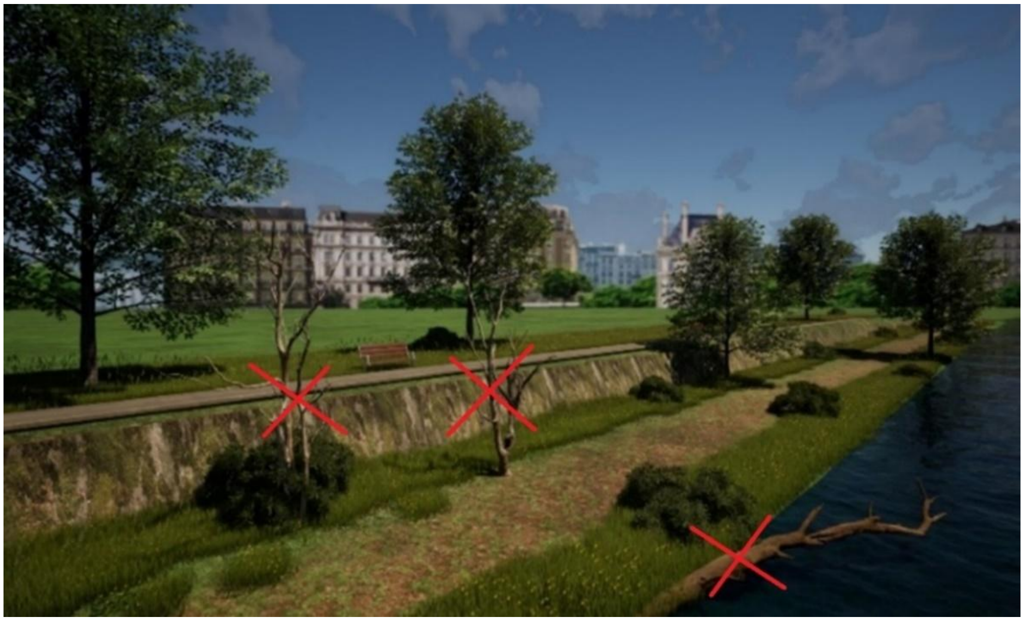

In our survey, the presence of dead wood in the riparian landscape can take three levels: none, low, medium. The levels surveyed correspond to the following landscape types:

Examples of riverside landscapes with dead wood

| | |
|---|---|
| 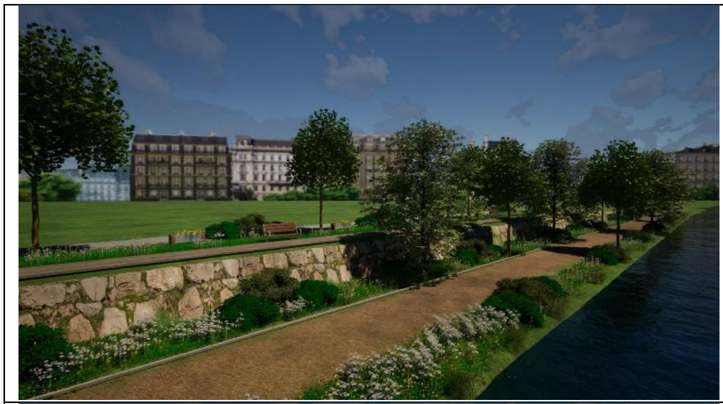 | **NONE**<br>No trees left to decay naturally |
| 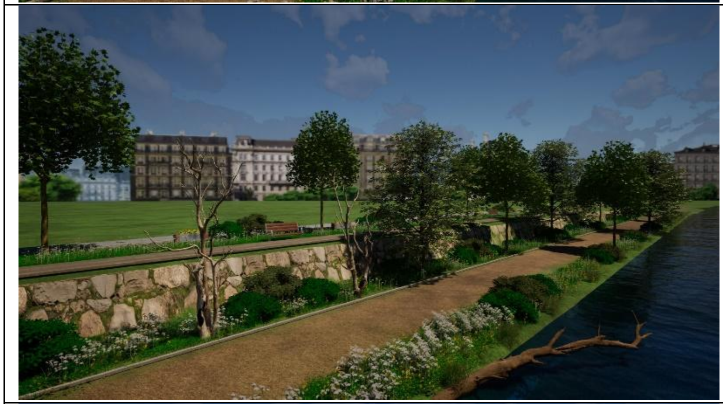 | **LOW**<br>Individual trees left to decay naturally |
| 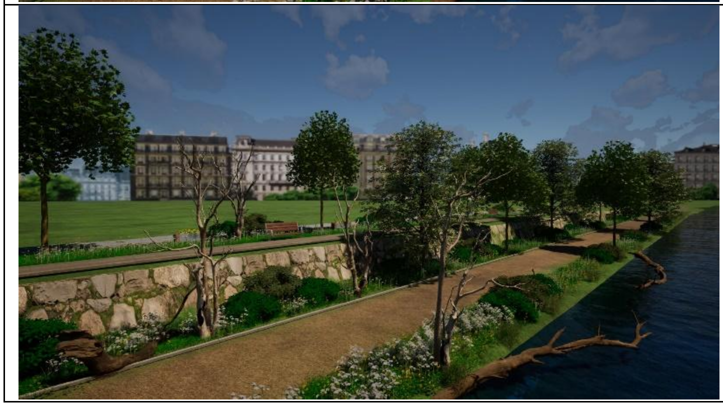 | **AVERAGE**<br>Average number of trees left to decay naturally |

**C7. Which of the landscapes shown do you find most attractive?**
**C8. Which of the landscapes shown do you find least attractive?**

**☐ Differences in the amount of dead wood do not matter to me**

**IMPERMEABLE SURFACES**

In urban areas, in many places natural permeable ground (e.g. soil, gravel, sand) is now being replaced by hardened impermeable surfaces such as concrete, asphalt or other water impermeable surfaces.

In our study, the proportion of impervious surface in a river valley can take three levels: **low, medium and high**.

**These levels correspond to the following landscape types**:

| | | |
|---|---|---|
| 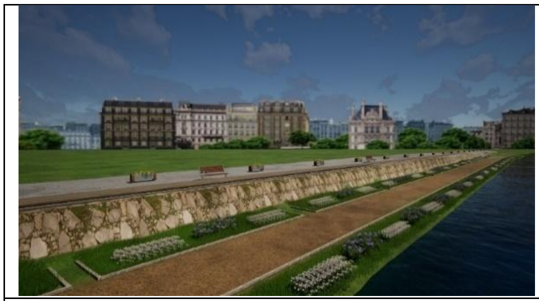 | 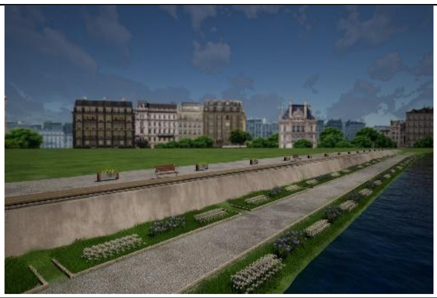 | 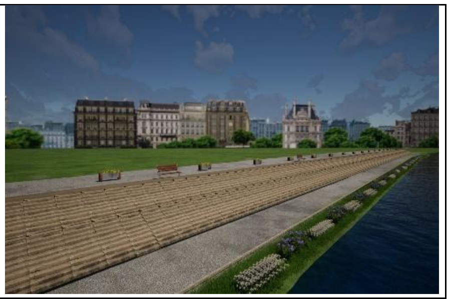 |
| **Low (10%)**<br><br>The proportion of impervious surface is **<u>10%</u>**, with the rest, i.e. 90%, being green areas | **Medium (40%)**<br><br>The proportion of impervious surface is **<u>40%</u>**, the rest i.e. 60% is green space | **High (90%)**<br><br>Share of impervious surface is **<u>90%</u>**, the rest i.e. 10% is green area |

**C9. Which of the landscapes shown do you find the most attractive?**
**C10. Which of the landscapes shown do you find least attractive?**

☐ **Differences in the proportion of impervious surfaces do not matter to me**

#### MANAGEMENT INTENSITY

The next characteristic we will consider is the intensity of management.  It describes the intensity and frequency of maintenance (e.g. planting, mowing, pruning).

In our study, management intensity takes three levels: **low, medium, high**.
These levels correspond to the following landscape types.

**Management intensity**
Example for low vegetation (grasses and herbs)

| | |
|---|---|
| 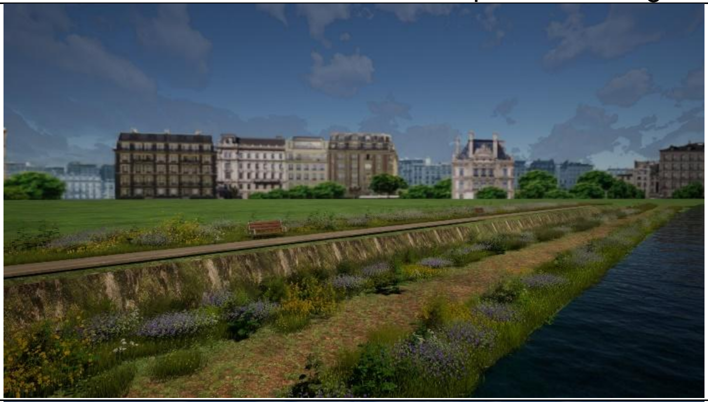 | **Low management intensity** (vegetation growing naturally, pruned and mown once a year) |
| 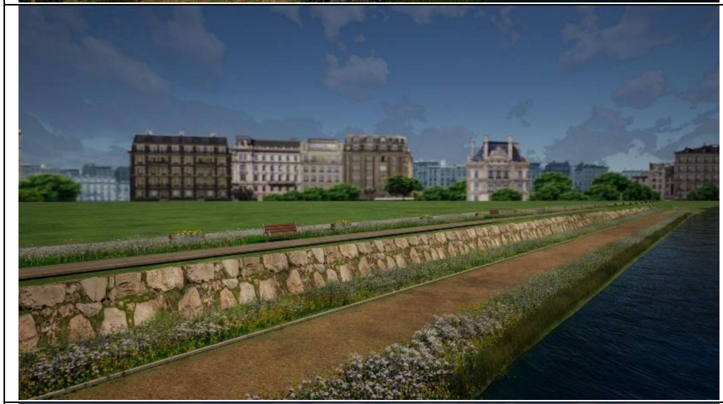 | **Medium management intensity** (vegetation partly planted and partly natural, pruned and mown twice a year) |
| 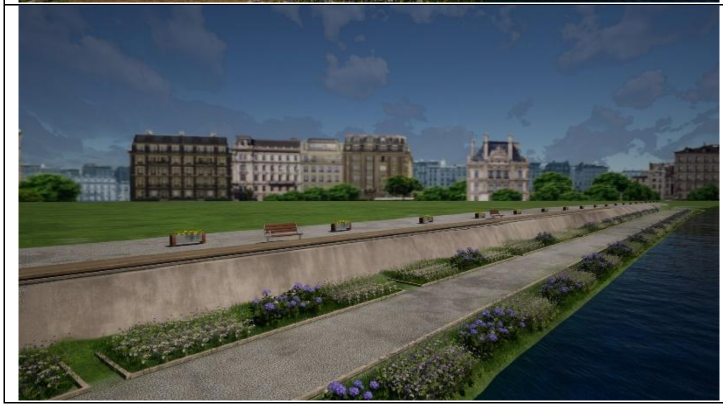 | **High intensity of management** (vegetation planted, pruned and mowed at least four times a year). |

**C11. Which of the landscapes shown do you find most attractive?**
**C12. Which of the landscapes shown do you find least attractive?**

☐ **Differences in management intensity do not matter to me**

**Management intensity**

Example for a landscape with medium vegetation (shrubs)

| | |
|---|---|
| 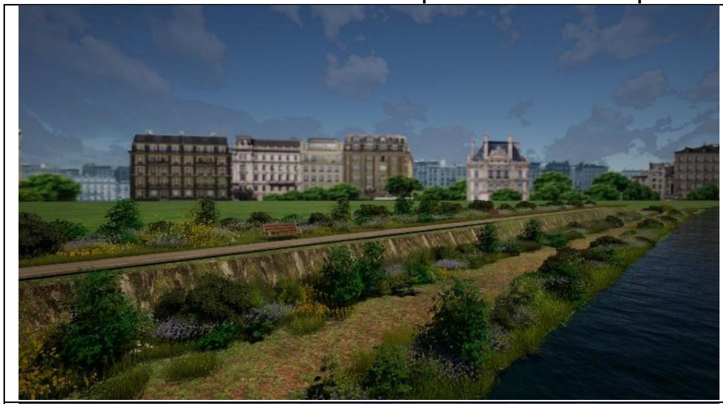 | **Low management intensity,** (vegetation growing naturally, pruned and mown once a year) |
| 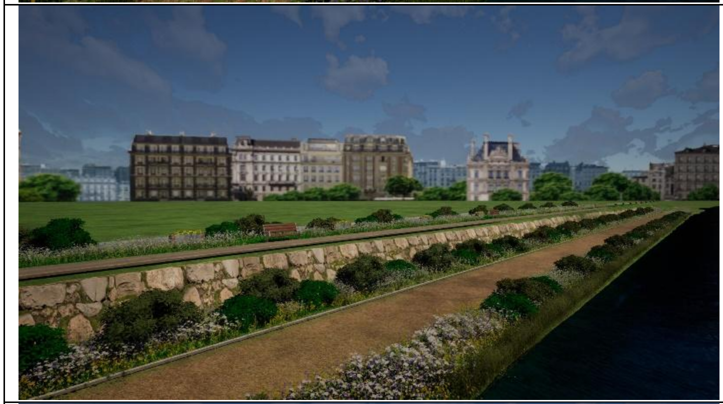 | **Medium management intensity,** (vegetation partly planted and partly natural, pruned and mown twice a year) |
| 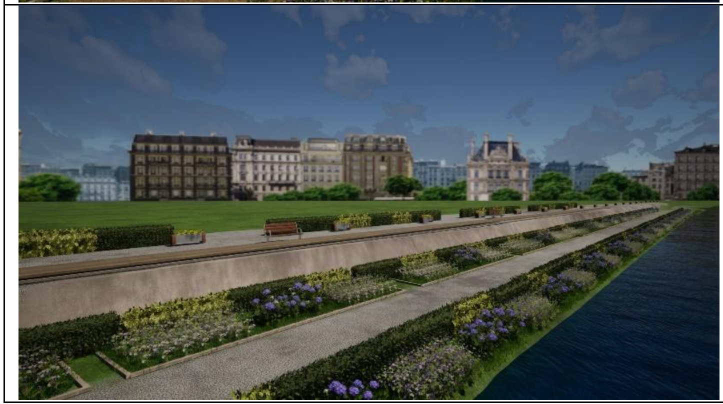 | **High management intensity,** (vegetation planted, pruned and mowed at least four times a year) |

**C11. Which of the landscapes shown do you find most attractive?**
**C12. Which of the landscapes shown do you find least attractive?**

**☐ Differences in management intensity do not matter to me**

**Management intensity**
Example for a landscape with tall vegetation (trees)

| | |
|---|---|
| 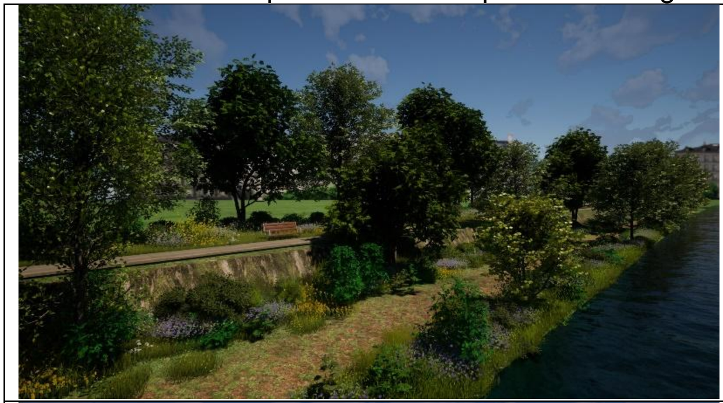 | **Low management intensity,** (vegetation growing naturally, pruned and mown once a year) |
| 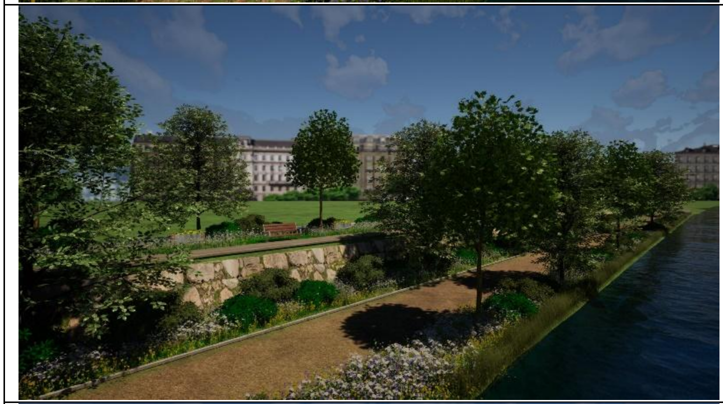 | **Medium management intensity,** (vegetation partly planted and partly natural, pruned and mown twice a year) |
| 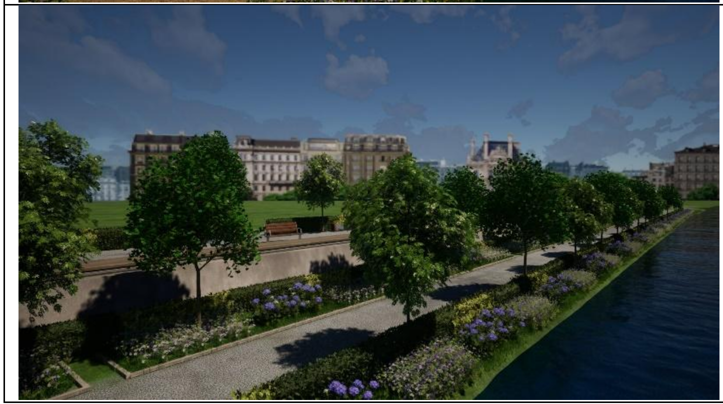 | **High management intensity,** (vegetation planted, pruned and mowed at least four times a year) |

**C11. Which of the landscapes shown do you find most attractive?**
**C12. Which of the landscapes shown do you find least attractive?**

☐ **Differences in management intensity do not matter to me**

**URBAN GREENERY MANAGEMENT CHANGE PROGRAMME**

In a moment, you will be shown 12 cards/screens presenting different approaches to the management of the riverside zone in your city. For each card, please indicate which scheme you think would be most beneficial for you. Three programmes will be shown on each card.

The first assumes a continuation of the current greenery management. If the current management of greenery in the riverside zone continues, the neighbourhood of rivers in many places in your city will, **in about 10 years' time**, be covered with highly simplified low vegetation and a high proportion of impervious surfaces. This landscape is schematically depicted in the visualisation below.

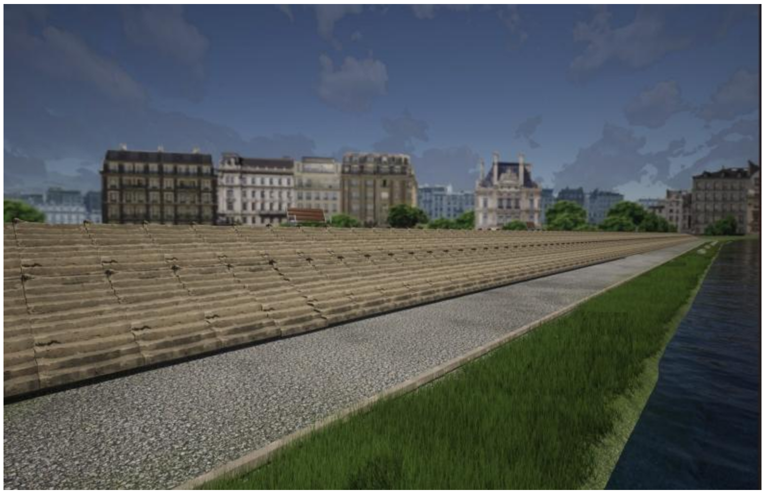

Two further programmes involve changes to the management of riverside areas in your city

Introducing changes in the management of urban greenery involves additional costs.
Therefore, one element of each programme is its cost.

Any of the options that involve making changes to the status quo will require additional costs for residents, including yourself, e.g. in the form of increased taxes. At present, the exact cost of introducing each of the schemes presented is unknown, but in making your decision, please treat the figures presented as the actual annual cost you would have to pay if the scheme were introduced.

Based on your opinion, as well as the opinion expressed by other residents, the local authority will be able to develop and implement an urban greenery management plan in a way that is in line with residents' expectations.

When filling in the questionnaire, please remember that your answers may influence the decisions on how the riverside areas in your city will be managed and what they will look like in the future.

**WHEN MAKING YOUR CHOICE, PLEASE REMEMBER THAT**

- Each combination should be considered independently of the others. In any case, please choose the scheme that you think is the best in terms of managing urban greenery.

- There are no right or wrong answers in the survey. Every opinion is important to us and can be used to develop future riverside green space management plans.

- Funding is needed to implement changes in landscape management. Such changes can sometimes come at a significant cost to local communities. Even supporting a low-intensity programme of green space management may involve the cost of removing existing infrastructure or the need to plant new green space.

- The exact costs of delivering the various scheme options are not yet known, so different cost options appear in the study.

**SECTION C - *Riparian vegetation valuation and management***

When analysing the potential cost of each scheme and making a selection, please also bear in mind that:

- In every household, money is needed for various other expenses.
- In each city there are many other investments for which additional funds can be used.
- If you think a particular option is too expensive for what it offers and you would not choose to pay that much for it, please do not choose it.
- On each card, one of the available options is the choice of "status quo" - i.e. no change, which does not incur any additional charges for you.

**//TREATMENT 1 STARTS WITH THE SELECTION TASKS//.**

| **TREATMENT 1 (50% of respondents)** | **TREATMENT 2 (50% of respondents)** |
|---|---|
| - no information | - providing additional information about services |

**// ONLY TREATMENT 2 //**

Please read the following information before choosing how to manage urban vegetation.

Areas of natural vegetation along rivers are important for nature. They provide suitable living conditions for many organisms, act as pathways for the movement of many animal species, and retain pollutants flowing into the river. These areas are also important for people, offering an attractive landscape and space for relaxation.

Unfortunately, many riverside areas are now under threat due to increasing urbanisation. Increasing pressure from urban expansion is leading to the destruction of riverside areas. In order to preserve their value and protect the organisms living there, it is necessary to protect the natural riverside vegetation to a greater extent than is currently the case.

In the chart below, we have outlined the main benefits provided by a riparian zone with well-preserved natural vegetation.

| |
|---|
| 1 - Recreation and leisure<br>Riverside areas are places for recreation, leisure and contact with nature. |
| 2 - Ecological corridors<br>Rivers with natural riparian vegetation are used by animals as routes that allow them to move. |
| 3 - Wildlife habitats<br>The proximity of water, lush vegetation, and the presence of trees, including decaying ones, provides suitable habitat for many species of animals and plants. |
| 4 - Temperature control<br>The presence of old trees provides shade, which lowers the water temperature. This is important for many aquatic organisms, especially in hot weather. |
| 5 - Dead wood<br>Fallen and standing hollow trees provide microhabitats for many species and mammals. Decaying wood is an important food source for numerous organisms. |

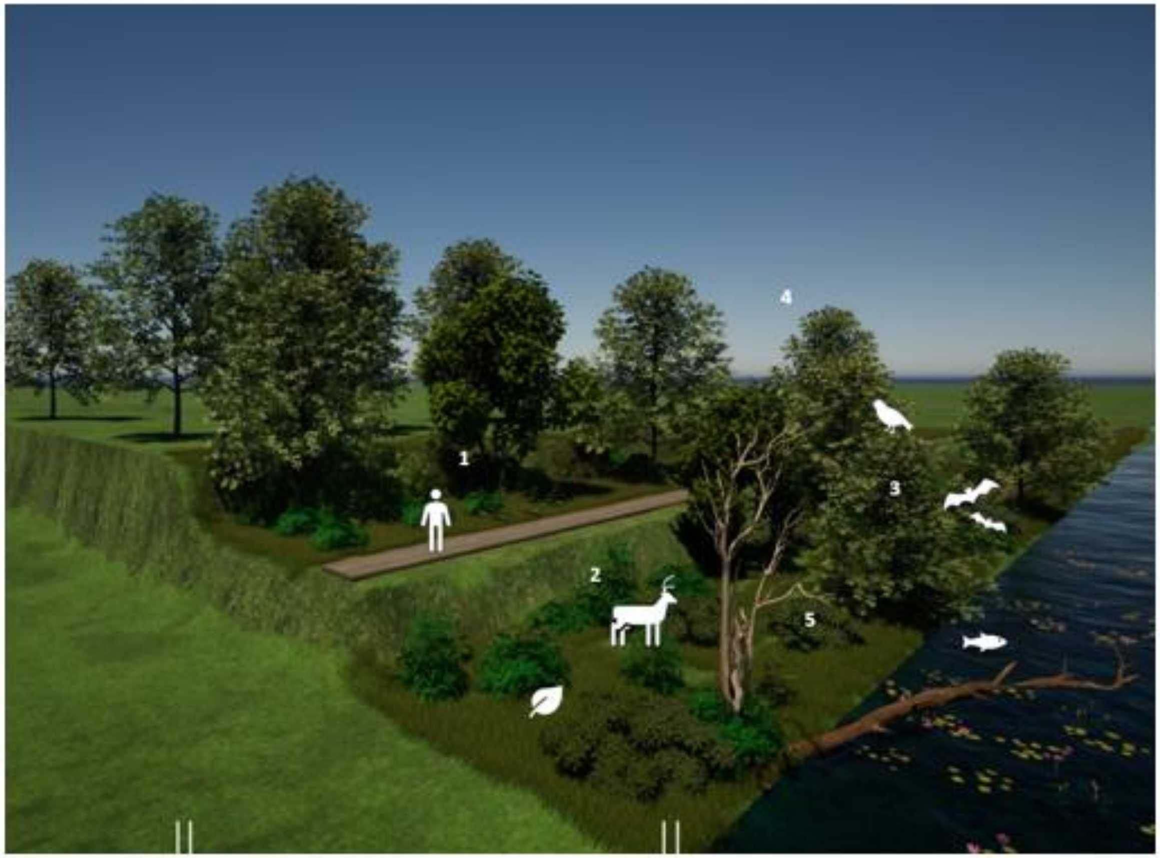

| Upland | Riparian | Stream/River |
|---|---|---|

Example selection sheet

| | Continuation of the current state | **Programme A** | **Programme B** |
|---|---|---|---|
| | 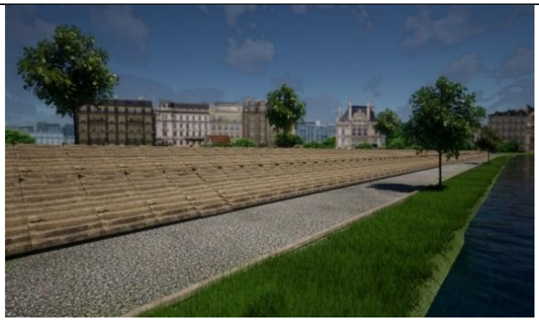 | 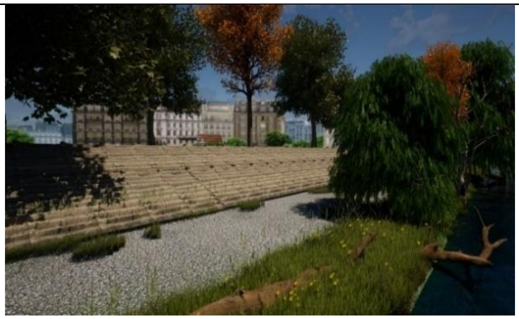 | 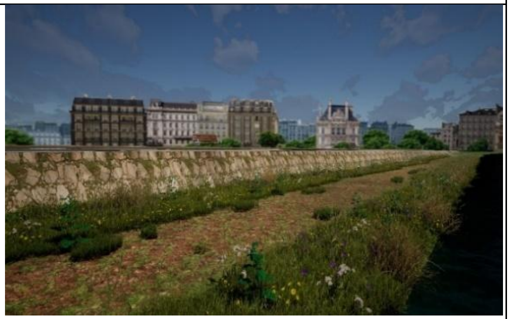 |
| Description of the highest vegetation storey (Type, Density, Species) | **Trees**<br>Low density of trees,<br>Low number of tree species | **Trees**<br>High tree density,<br>High number of tree species | **Grasses, herbs**<br>High density of grasses and herbs,<br>High number of species of grasses and herbs |
| Presence of dry wood | Absence of | Small amount | None |
| Intensity Maintenance of vegetation | High | Medium | Low |
| Impervious surface | 70% of the surface | 70% of surface | No concrete |
| **Cost For you** | **0 zł/year** | **25 zł/year** | **50 zł/year** |
| Your choice | ☐ | ☐ | ☐ |

Treatment B: When making your choices, please remember that well-preserved natural riparian vegetation provides many benefits in addition to landscape value (Active link to earlier sheet)

**SECTION G - *Motivation***

**SHOW QUESTION G1 ONLY TO RESPONDENTS WHO ANSWERED "NO CHANGE SCENARIO" (STATUS QUO) TO ALL CHOICE TASKS.**

G1. In all choice situations you indicated a continuation of the current management of the riverside greenery. Please indicate which of the following statements best describes your motives for this behaviour. Please select up to 2 answers from the list below

| |
|---|
| Understanding the schemes was difficult. Choosing the "no change scenario" was the easiest option |
| I would not like my tax money to be spent on urban greenery in my city |
| How the greenery looks in the neighbourhood of the river in my city does not matter to me |
| All programmes, with the exception of the "No change" scenario were too expensive |
| Improving the management of urban greenery should be financed using other means, not my taxes |
| The taxes I currently pay should be enough to fund urban green space management |

Other, which ones? _______________________________

**SHOW THE FOLLOWING QUESTIONS TO ALL RESPONDENTS, REGARDLESS OF THEIR CHOICE.**

G4. To what extent, do you agree or disagree with the following statements? (please choose)

| | Totally agree | Rather agree | I rather disagree | I completel y disagree | I do not know |
|---|---|---|---|---|---|
| The new programme should focus on improving the attractiveness of riparian areas for people | | | | | |
| The new programme should put more emphasis on protecting biodiversity than on improving the attractiveness for people | | | | | |
| The new programme should put more emphasis on improving the attractiveness of riverine areas for people than on protecting biodiversity | | | | | |
| Biodiversity protection and human wellbeing should be given equal consideration in the implementation of new programmes in the riverine area | | | | | |

G5. Please indicate to what extent you think the results of this survey will be used to change the management of riverside areas in your city. (1 answer maximum)

| |
|---|
| I am strongly convinced that they will not have any impact |
| I am convinced they will have no impact |
| I assume they are unlikely to have an impact |
| I assume they might have some influence |
| I am convinced they will have an impact |
| I am strongly convinced they will have an impact |
| I don't know, I have no opinion |

G6. Please indicate the extent to which you think the results of this survey will be used to determine future changes in the amount of taxes paid to change the management of riparian areas in your town. (1 answer maximum)

| |
|---|
| I am strongly convinced that they will not have any impact |
| I am convinced they will have no impact |
| I assume they are unlikely to have an impact |
| I assume they might have some influence |
| I am convinced they will have an impact |
| I am strongly convinced they will have an impact |
| I don't know, I have no opinion |

**SECTION H -** ***Concluding questions***

**H1. Have you ever heard of the term biodiversity?**
Options:
a) I have heard of it and I know what it means;
b) I have heard of it but do not know what it means;
c) I have never heard of it;
d) I don't know

**PLEASE READ:**
**Biodiversity - or biodiversity - is the term for the variety of life on Earth (e.g. plants, animals, genes, but also ecosystems such as forests, oceans, etc.) of which we are an integral part. Biodiversity in Europe and other parts of the world is being lost and degraded due to human activities.**

**H2. Please indicate to what extent you agree or disagree with the following statements**

| | **Totally agree** | **Rather agree** | **I rather disagree** | **Complet ely disagree** | **I do not know** |
|---|---|---|---|---|---|
| We have a duty to care for nature | | | | | |
| Our health and well-being are based on nature and biodiversity | | | | | |
| Biodiversity is important for our long-term economic development | | | | | |
| We have a responsibility to prevent further loss of biodiversity | | | | | |

**H3. Please indicate which of the following actions you think would be most important for biodiversity conservation. (Max 5 answers) maybe 3?**

| |
|---|
| Expanding areas where nature is protected |
| Strengthening existing nature conservation and biodiversity protection rules |
| Better implementation of existing nature and biodiversity conservation rules |
| Allocating more funding to nature and biodiversity conservation |
| Promoting research into the impacts of biodiversity loss |
| Better educating and informing citizens about the importance of biodiversity |
| Taking biodiversity into account when planning new infrastructure |
| Implementation of new and innovative nature conservation strategies |
| Promoting and protecting urban biodiversity |
| Other, which? |
| None |
| Don't know |

**Do you have any comments or feedback that you would like to share with us?**

Please click on the "Send**"** button to complete the survey.

**By clicking "Submit", you declare that you have voluntarily taken part in this scientific study and agree that your anonymous information will be used for the specific purposes of this scientific study.**

#### Thank you very much for your time and participation in the study!
